# scHG: a supercell framework with high-order graph learning enables hyper-fast multi-omics analysis

**DOI:** 10.64898/2025.12.19.695346

**Authors:** Yixiang Huang, Yuan Gan, Xinqi Gong

## Abstract

Multi-omics profiling—spanning proteomics, transcriptomics, and additional omics data types—is rapidly advancing, providing increasingly detailed maps of cellular identity and function. In parallel, computational methods have evolved to integrate diverse omics with increasing precision in resolving cellular heterogeneity and delineating subtle subpopulation structures. Yet, capturing rare cell population while maintaining computational tractability remains a major challenge. Here, we introduce the supercell paradigm, in which expression-coherent cells are compressed into candidate units for rare cell population. Supercells are constructed using angle-aware similarity metrics and second-order co-occurrence neighbors, with impurity cells pruned by degree centrality. To address scalability, we implement sparse matrix optimization and iterative high-order graph updates, enabling efficient integration of large-scale multi-omics datasets. Building on this framework, we develop scHG, a high-order graph learning approach guided by an omics-weighted optimizer that adaptively balances contributions from gene expression, surface proteins, and chromatin accessibility. Across six benchmark datasets (up to 30672 cells), scHG consistently outperforms state-of-the-art methods, improving mean ARI and NMI by 3.97% and 3.54%, respectively, while reducing runtime by 26.40%. Beyond performance gains, scHG was able to resolve fine-grained cellular heterogeneity within conventionally defined T cell populations, distinguishing subpopulations such as memory-like or progenitor-exhausted T cells and innate-like cytotoxic T cells. At the same time, the supercell framework uncovered rare populations, including dendritic-cell populations and NK-like B cells, that remained masked at the cluster level under standard pipelines. These results underscore both the rare-cell detection capability and the computational efficiency of scHG. Our code and data are available at http://mialab.ruc.edu.cn/scHG_code/zip.

## Introduction

Thanks to the rapid development of bioinformatics tools, multi-omics analysis has flourished in recent years. As a fundamental task, clustering provides the foundation for a wide range of downstream analyses, including cellular heterogeneity characterization, analysis of cell development trajectory, and cell-cell communication inference. Given that different omics modalities—such as the proteomics, transcriptomics, and chromatin accessibility—capture complementary aspects of cellular states, an essential challenge is how to effectively integrate these heterogeneous datasets for joint clustering. This integration problem is naturally formulated as a multi-view clustering task, where each omics modality represents a distinct view of the same underlying biological system.

Multi-view clustering has been extensively explored in the machine learning community through a diverse set of approaches. A seminal work by Bickel and Scheffer [1] introduced one of the earliest probabilistic frameworks to capture shared latent structures across views, laying the groundwork for future studies. Follow-up methods included orthogonalization-based representation learning [2], canonical correlation analysis [3], scalable extensions of K-means [4], and structured sparsity-based clustering and feature selection [5]. More recent advances such as PMSC [6] and SMSC [7] have incorporated subspace and spectral embedding techniques, while models like GCFAgg [8] and auto-weighted multi-view clustering [9] emphasize feature aggregation and adaptive weighting for handling heterogeneous data sources.

These general machine learning techniques have profoundly shaped the early development of bioinformatics-specific multi-omics clustering algorithms. Zhang et al. proposed an integrative strategy to uncover multi-dimensional functional groups from cancer genomic datasets, enabling the discovery of biologically coherent patterns across modalities [10]. Wu et al. developed a low-rank approximation framework that accelerated dimension reduction and improved cancer molecular subtyping by efficiently handling large-scale heterogeneous data [11]. Yang and Michailidis advanced this line of work by introducing a non-negative matrix factorization approach tailored for heterogeneous multi-modal omics, capable of detecting coordinated structures across diverse data types [12]. Subsequent efforts such as scAI [13] utilized non-negative matrix factorization to jointly embed transcriptomic and epigenomic data, while LIGER [14] performed integrative decomposition into shared and dataset-specific factors. BREM-SC [15] adopted a Bayesian random effects mixture model to account for inter-cell and inter-omic variability, and HONMF [16] incorporated hypergraph-based regularization into matrix factorization. More recently, scMNMF and GSTRPCA [17, 18] extend matxrix factorization and robust PCA, respectively, to capture biological structures across omics layers. Together, these non-graph-based methods established an essential foundation for multi-omics integration. They not only improved clustering accuracy but also enabled downstream applications such as differential expression analysis, batch effect correction, and the reconstruction of cell developmental trajectories.

In recent years, graph-based approaches have emerged as a competitive paradigm for multi-view clustering, particularly suited to the complexity and noise inherent in omics data. By leveraging relational information both within and across views, these methods mitigate the shortcomings of earlier factorization-based techniques, offering improved structural fidelity and resilience to outliers. Representative advances include GBS [19], which established a general graph-based framework for multi-view learning, and CGD [20], which employed cross-view graph diffusion to strengthen inter-view communication. Methods such as GMC [21] and Consensus Graph Learning [22] further emphasized the construction of unified or consensus graphs that distill view-shared structures. In the bioinformatics setting, Jiang et al. [23] introduced a Laplacian optimization framework for robust clustering of multi-omics data, demonstrating the adaptability of graph-based formulations for encoding feature dependencies. These graph-based methods exhibit *O*(*n*^3^) complexity, primarily due to the eigen-decomposition required in spectral clustering [19, 21–23] and the large-scale matrix multiplications involved [20]. Taken together, these developments consolidate graph-based models as a robust and versatile framework for multi-view integration and clustering across both machine learning and biological domains.

However, despite their partial effectiveness, current graph-based methods still face two major challenges that limit their scalability and resolution on large-scale multi-omics datasets. First, since eigen-decomposition in spectral clustering or large-scale matrix multiplications, graph-based methods generally incur a computational cost of *O*(*n*^3^), making them infeasible for large-scale omics datasets. Second, rare cell populations remain difficult to detect, since separating small yet expression-coherent groups without disrupting global cluster integrity requires capturing subtle higher-order relationships. This calls for further exploration of distance metrics and refined representations of cell-to-cell similarity that go beyond traditional pairwise measures. Together, these challenges underscore the need for frameworks that can efficiently incorporate higher-order structure while maintaining computational tractability.

To overcome these limitations, we propose a novel high-order neighbor-aware coarse-grained multi-omics graph clustering framework. The core innovation of our approach lies in the introduction of the supercell concept, which aggregates cells into biologically meaningful units when their expression profiles display higher-order consistency that is stably preserved across multiple omics layers (Fig. 1). These units operate at an intermediate resolution between single cells and tissue-scale structures, providing a principled means to balance fine-grained heterogeneity with computational efficiency. The formulation of supercells serves two primary objectives. First, by aggregating cells into coherent units, we dramatically reduce the effective sample size, thereby enhancing scalability and rendering our framework suitable for large-scale datasets. Second, supercells enable the capture of more subtle cellular subtypes without increasing the number of clusters, thereby facilitating the identification of rare cell populations. Conceptually, our notion of supercells draws inspiration from earlier coarse-grained strategies such as the metacell representation in MetaCell [24], the pseudobulk paradigm used for robust differential testing in Squair [25], and the cellular neighborhood framework for spatially organized units in Sch”urch [26]. More recent approaches, including SEACells [27] and PAGA [28], highlight the utility of intermediate-resolution structures for trajectory inference and state discovery. Unlike these methods, however, our supercell framework explicitly integrates multi-omics concordance and high-order neighborhood information, enabling a unified treatment of scalability, rare-cell recognition, and multi-omics integration.

**Fig 1.**
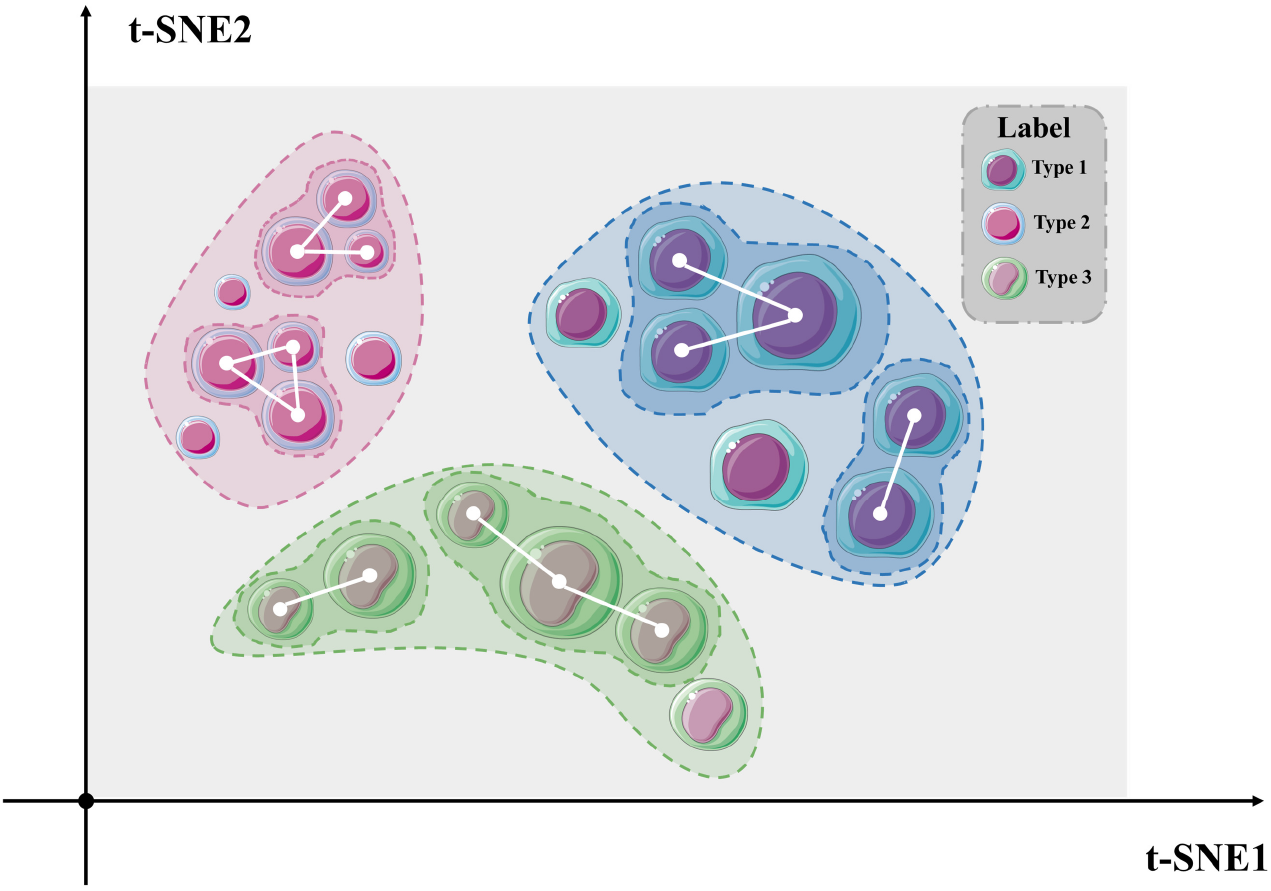
Visualization of t-SNE based on supercell clustering results.

Our study makes three key contributions. First, we validate the effectiveness of the supercell concept in multi-omics integration for the first time, showing on the PBMC10× benchmark that it reveals rare dendritic-cell populations and NK-like B cells otherwise masked at the cluster level. Second, building on supercells, we develop a scalable graph-based fast multi-omics clustering framework that consistently surpasses state-of-the-art methods, achieving mean gains of 3.97% in ARI and 3.54% in NMI, while clustering datasets of over 30000 cells in under 20 minutes^1^. Finally, we propose a high-order graph model that combines second-order co-occurrence neighbors with angle-aware metrics and stochastic pruning, delivering a 14% average improvement over baselines and adaptively determining cluster numbers without prior assumptions.

## Results

### Overview of scHG

Our workflow delivers a two-stage, graph-based partitioning that first groups cells into supercells and then organises those supercells into higher-level clusters (Fig. 2).

**Fig 2.**
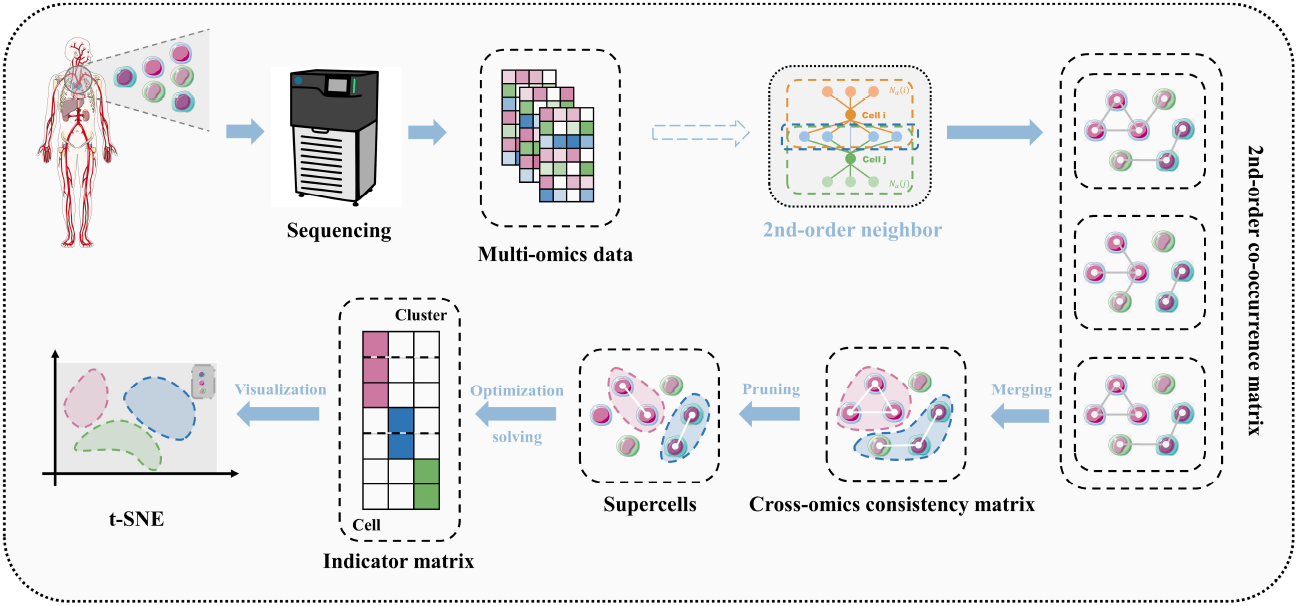
The architecture of scHG. Our integrated workflow comprises: (1) second-order co-occurrence neighbor extraction from input multi-omics data; (2) cross-omics consistency fusion; (3) probabilistic pruning to identify high-order structural units (supercells); (4) supercells clustering via optimization algorithms; and (5) visualization with downstream analytical outputs.

#### Stage 1: Supercell formation

For each omics layer, we build a *γ*-nearest-neighbor graph with Pearson-correlation weights, promote it to a second-order graph by requiring mutual *α* - neighborhood and at least *β* shared neighbors, and then fuse all layers into a cross-omics consistency graph. Connected components in this fused graph form supercell candidates; degree - centrality – based pruning removes low-coherence outliers, yielding biologically coherent supercells that retain both local co-expression and global manifold structure.

#### Stage 2: Supercell clustering

The final optimized model formulation is given by Eq. (1) (see Section for derivation):

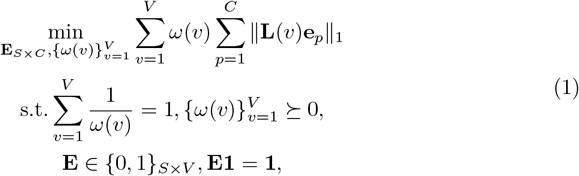

where **E**_*S*×*V*_ = (*e*_1_, *e*_2_, …, *e*_*V*_) denotes the clustering result of supercells, 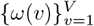 represents the optimization weights for distinct omics layers, **L**(*v*) is the Laplacian matrix of **AS**(*v*) = (*as*_*yz*_(*v*)) defined in Eq. (16).

We iteratively optimise cluster labels and modality weights with block-coordinate descent: (i) compute a Laplacian matrix for each omics-specific supercell graph and update a closed-form weight that down-weights poorly cluster-separated modalities, (ii) build a composite Laplacian matrix and place each supercell in the cluster that yields the lowest inter-cluster similarity. Alternating these steps converges to a modality-weighted partition that maximises between-cluster separation while preserving cross-omic consensus. A more detailed description of the workflow is provided in the section.

### scHG enhances the resolution of cell subpopulation recognition

For the clustering results obtained by scHG on the PBMC10× dataset, we performed one-sided Welch’s *t*-tests^2^ to identify enriched markers across clusters and subsequently assigned cell-type labels (Table 1, Supplementary Table 1). Notably, Clusters 3 and 7 shared a core set of T-cell markers—CD3, CD4, CD25, CD45RO, PD-1 and CD127—indicating that both represent CD4+ T-cell populations.

**Table 1.** Top enriched markers in cluster 3 and cluster 7 based on one-sided Welch’s t-test.

| Cluster | Marker | Type | p-value |
| --- | --- | --- | --- |
| Cluster 3 | CD3_TotalSeqB | Protein | $< 1 \times 10^{-300}$ |
| | CD4_TotalSeqB | Protein | $< 1 \times 10^{-300}$ |
| | CD127_TotalSeqB | Protein | $2.27 \times 10^{-204}$ |
| | CD45RO_TotalSeqB | Protein | $3.14 \times 10^{-58}$ |
| | PD-1_TotalSeqB | Protein | $1.41 \times 10^{-33}$ |
| | CD25_TotalSeqB | Protein | $2.95 \times 10^{-32}$ |
| | CD27 | RNA | $8.48 \times 10^{-67}$ |
| | HSPA8 | RNA | $1.26 \times 10^{-23}$ |
| | AQP3 | RNA | $1.04 \times 10^{-21}$ |
| | PIM2 | RNA | $2.36 \times 10^{-18}$ |
| Cluster 7 | CD16_TotalSeqB | Protein | $9.87 \times 10^{-21}$ |
| | CD4_TotalSeqB | Protein | $3.79 \times 10^{-17}$ |
| | CD15_TotalSeqB | Protein | $1.44 \times 10^{-14}$ |
| | CD45RO_TotalSeqB | Protein | $3.15 \times 10^{-9}$ |
| | CD3_TotalSeqB | Protein | $1.50 \times 10^{-8}$ |
| | CD127_TotalSeqB | Protein | $1.61 \times 10^{-6}$ |
| | CD25_TotalSeqB | Protein | $2.66 \times 10^{-4}$ |
| | PD-1_TotalSeqB | Protein | $6.71 \times 10^{-4}$ |
| | TIGIT_TotalSeqB | Protein | $3.38 \times 10^{-2}$ |
| | CCND2 | RNA | $2.11 \times 10^{-3}$ |

Beyond this common signature, however, each cluster exhibited distinct molecular programs. Cluster 3 was distinguished by transcriptional enrichment of activation- and proliferation-related genes, including *CD27, PIM2* and *HSPA8*, together with a broad set of cell-cycle regulators such as *MKI67, CCNB1, EZH2, CHEK1* and members of the *MCM* family. This molecular profile points to a stem-like or central-memory T-cell subpopulation with features of early exhaustion, consistent with previous observations [29, 30].

By contrast, Cluster 7 displayed a markedly distinct surface phenotype, with significant enrichment of CD16, CD15 and TIGIT in addition to the shared T-cell markers. The co-expression of CD16 and CD15—typically associated with NK cells and granulocytes, respectively—points to the emergence of an innate-like cytotoxic T-cell subpopulation. Such CD16^+^ T cells have been recognized as functionally distinct subpopulations with heightened cytolytic potential [31, 32], while CD15 expression, though rare, has been observed in activated *γδ* T cells and NKT-cell subpopulations [33].

Taken together, these results demonstrate that although both clusters fall within the CD4+ T-cell population, scHG distinguishes subpopulations with divergent functional programmes: Cluster 3 most likely represents memory-like or progenitor-exhausted T cells, whereas Cluster 7 exhibits hallmarks of innate-like cytotoxic T cells. These observations underscore the capacity of our framework to enhance the resolution of cell subpopulation identification within noisy, high-dimensional multi-omics datasets, independent of any a priori label information.

### Label-free identification of clinically significant B-cell biomarkers

To assess the biological validity of our framework, we examined Cluster 5 (Supplementary Table 1), which is labelled as a B-cell population by the reference annotations. Applying a one-sided Welch’s *t*-test, we identified a panel of markers that are significantly enriched in both the transcriptomic and proteomic modalities.

At the protein modality, classical B-cell surface markers— CD19 (*p* = 2.03 × 10^−168^) and CD45RA (*p* = 2.47 × 10^−118^)—were markedly enriched in Cluster 5 [34]. At the transcriptomic modality, we observed significant up-regulation of canonical B-cell genes, including *CD74* (*p* = 1.43 ×10^−158^) [35], *MS4A1* (*p* = 4.08 × 10^−104^) [36], *BANK1* (*p* = 5.89 × 10^−101^) [37, 38], and *CD79A* [39]. Moreover, multiple MHC class II genes—*HLA*-*DQB1, HLA*-*DQA1*, and *HLA*-*DRB1* —were strongly over-expressed (*p <* 1 × 10^−50^) [40], collectively corroborating the B-cell identity of this cluster.

These findings demonstrate that our framework reliably recovers biologically meaningful signatures congruent with established cell identities. The concordant enrichment of B-cell markers at both the protein and transcriptomic modalities in Cluster 5 highlights the robustness and accuracy of the proposed algorithm.

Importantly, these markers are not only canonical B-cell identifiers but also represent validated or emerging therapeutic targets. For instance, CD19 is one of the most prominent B-cell antigens exploited in contemporary immunotherapy, underpinning multiple CAR-T products and monoclonal antibodies [41]. *MS4A1* constitutes the molecular target of rituximab and next-generation anti-CD20 antibodies, which are widely deployed in the treatment of B-cell lymphomas and autoimmune diseases [36]. *CD79A*, an essential component of the B-cell receptor (BCR) complex, has likewise been proposed as a druggable node for suppressing aberrant BCR signalling [42]. Finally, *CD74*, which functions in MHC class II antigen presentation and serves as a receptor for macrophage migration inhibitory factor (MIF), is currently being pursued with small-molecule inhibitors in pre-clinical studies [43].

In the aggregate, these findings demonstrate that—even in the absence of prior cell-type annotations—our algorithm can uncover biologically and clinically actionable markers, thereby highlighting its applicability to both fundamental immunology research and translational drug discovery.

### Cross-omics enrichment validates scHG in uncovering coordinated NK-cell effector programs

To further elucidate the functional implications of the markers up-regulated in the clusters identified by scHG, we conducted Gene Ontology (GO) enrichment analysis at both the transcriptomic and proteomic levels (Supplementary Table 2).

We analysed Cluster 2, which is annotated as a natural-killer (NK) cell population. At the transcriptomic level, as depicted in Fig. 3, the bar plot (Fig. 3d) reveals that the most significantly enriched Gene Ontology (GO) terms—including *leukocyte-mediated immunity, cell killing*, and *natural killer cell mediated cytotoxicity* —mirror the cytotoxic immune function of this cluster. The bubble plot (Fig. 3c) shows that pathways related to NK-cell activation, immune regulation, and degranulation score highest, with terms such as *cytolytic granule* achieving the greatest statistical significance. The circular plot (Fig. 3a) contrasts the total and differentially expressed genes (DEGs) associated with each GO term, underscoring the pronounced enrichment in immune-related pathways. Finally, the chord diagram (Fig. 3b) maps key up-regulated genes to representative biological processes, showing that *GZMB, NKG7*, and several *KLR* family members are pivotal mediators of cell-killing and NK-cell immune responses.

**Fig 3.**
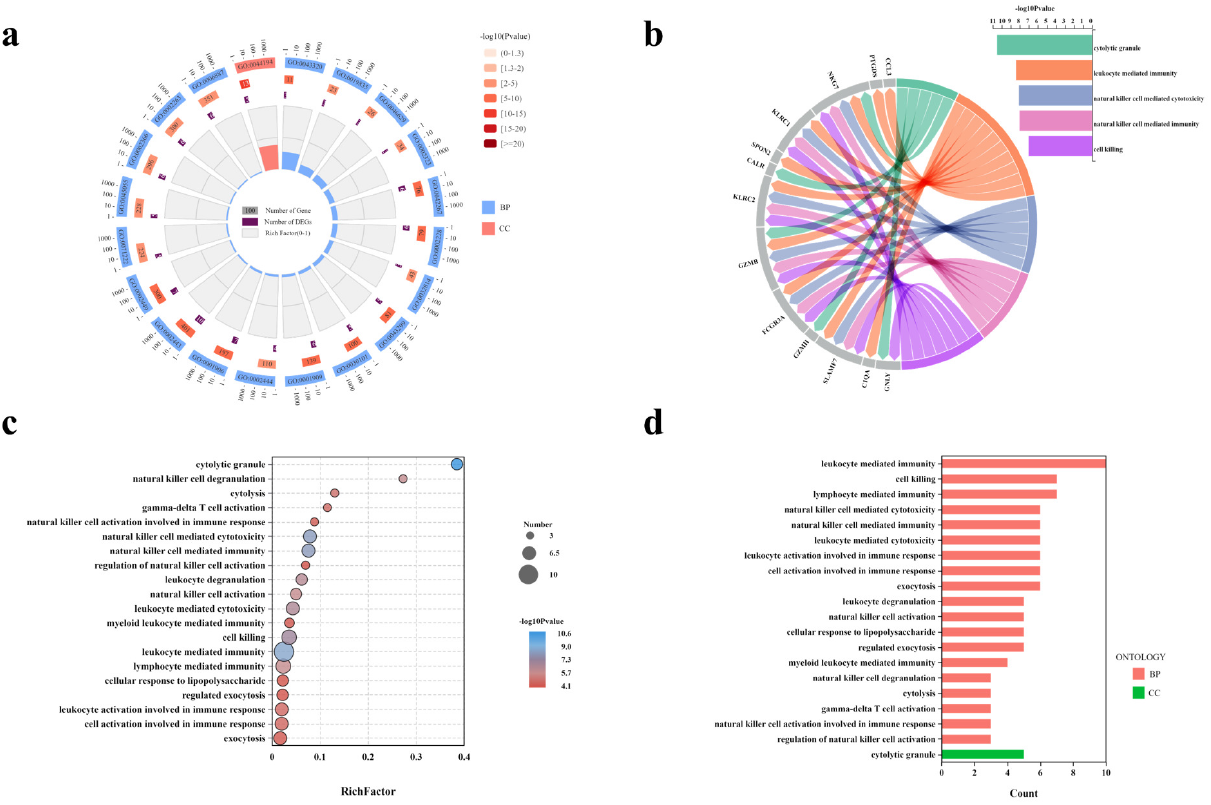
Functional enrichment and gene-pathway association analysis of upregulated genes. (a) GO term circular plot showing the number of total and differentially expressed genes (DEGs) associated with each GO term, categorized into biological processes (BP) and cellular components (CC). (b) Chord diagram linking representative upregulated genes to their enriched immune-related biological processes, emphasizing genes involved in cytolytic granule formation and natural killer cell-mediated cytotoxicity. (c) GO enrichment bubble plot of upregulated genes, displaying significantly enriched biological processes related to immune cytotoxicity and NK cell activity. (d) Bar plot of GO terms significantly enriched among upregulated genes.

At the proteomic level, we observe enrichment patterns that mirror and sharpen the transcriptomic signal (Fig. 4). The bar plot in the lower-right quadrant (Fig. 4d) ranks the twenty most significant Gene Ontology terms and shows three biological-process categories in clear prominence: “external side of plasma membrane”, “positive regulation of tumour necrosis factor production” and “signalling receptor binding”. Together, these terms indicate that the differentially abundant proteins are concentrated at the cell surface, are actively engaged in driving TNF-mediated cytokine release and operate as key receptor–ligand hubs that coordinate NK-cell effector responses. The bubble plot (Fig. 4c) directly above integrates effect size and significance, revealing that terms such as “positive regulation of tumor necrosis factor production” and “external side of plasma membrane” combine a high Rich Factor (0.02 and 0.015, respectively), large protein counts and strong statistics (-log_10_*p* values close to 9). Together they point to surface-localised molecules that not only mediate antibody-dependent interactions but also potentiate TNF-driven cytokine release, thereby delineating the membrane-centred signals that prime NK-cell activation. A more granular perspective is provided by the circular GO diagram in Fig. 4a, which arranges enriched terms around the three GO ontologies, encoding for each term its protein count, regulation direction and Rich-Factor scale. In the biological-process (BP) sector (peach), seven terms stand out: *positive regulation of interferon-γ production* (GO:0032819), *Fc-gamma-receptor signalling pathway* (GO:0038094), *regulation of T-cell-receptor signalling* (GO:0050856), *negative chemotaxis* (GO:0001960), *regulation of phagocytosis* (GO:0050764), *organelle inheritance* (GO:0048304) and *positive regulation of natural-killer-cell-mediated cytotoxicity* (GO:0045059). Together they depict an activated immune state that integrates cytokine release, antibody-dependent triggering, migratory cues and effector NK-cell functions. The cellular-component (CC) ring (violet) is dominated by membrane-associated terms—GO:0009897 (external side of plasma membrane) and GO:0032059 (bleb)—highlighting that many differentially abundant proteins localise to the cell surface or dynamic membrane protrusions. Although fewer in number, the blue molecular-function (MF) segments are highly significant; they include *signalling-receptor binding* (GO:0005102), *transmembrane receptor protein tyrosine-phosphatase activity* (GO:0005001) and *heparan-sulfate proteoglycan binding* (GO:0043395), capturing the specific ligand–receptor chemistries and enzymatic activities that modulate NK-cell signalling thresholds. Altogether, the distribution of GO terms confirms that the proteomic signature of our marker set is concentrated in pathways governing immune activation at the cell surface, chemotactic positioning and NK-cell effector mechanisms. Finally, the network panel (Fig. 4b) maps representative proteins—P08637, P08575 and Q495A1—to their top-ranking GO terms and shows that these hubs converge on pathways such as Fc-gamma receptor signalling, TNF induction and NK-cell proliferation, underlining their coordinated contribution to cytotoxic effector function. Taken together, the four panels demonstrate a coherent proteomic programme centred on antigen-dependent cytotoxicity, membrane localisation and checkpoint regulation, thereby corroborating the cross-omics consistency and biological validity of the marker set identified by our framework.

**Fig 4.**
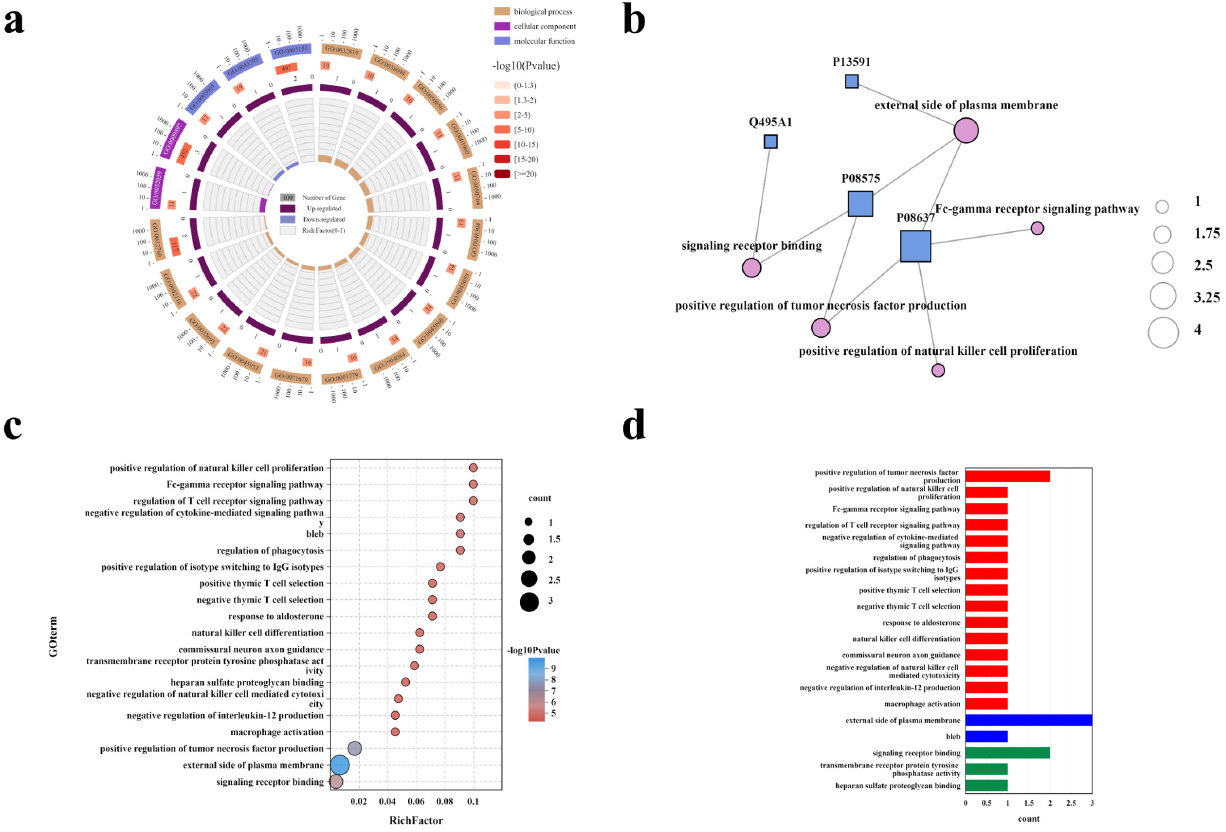
GO enrichment analysis of upregulated protein-level markers reveals key immunoregulatory pathways and molecular interactions. (a) Circular GO plot displaying the number and direction of differentially expressed proteins in each enriched term, with category annotations and enrichment scores. (b) Network plot linking representative proteins (by UniProt ID) to key enriched GO terms, highlighting function-specific interactions. (c) Bubble plot showing the top GO terms enriched by protein-level markers. Size reflects gene count, color indicates p-value, and x-axis denotes Rich Factor. (d) Bar plot of significantly enriched GO terms based on protein-level markers.

### Supercell reveals rare cell populations

The supercell framework aggregates cells with coherent expression profiles into compact units. Supercells comprising a large number of cells are more likely to display distinct expression patterns relative to the surrounding cluster, thereby facilitating the identification of rare cell populations. We illustrate this principle with two examples.

In the PBMC10× dataset, one-sided Welch’s *t*-tests revealed that Cluster 6 was enriched for CD3, CD8A, CD8B, GZMK/GZMH/NKG7, CD45RO, PD-1, TIGIT, and CD127, consistent with a CD8^+^ T-cell phenotype. Intriguingly, several dendritic-cell markers—*CD1C, CD1E, CLEC10A, FCER1A*, and *CD74* —were also detected, suggesting the presence of an antigen-presenting cell subset. To disentangle this heterogeneity, we separately analysed Supercell 62—the largest unit within Cluster 6—and Cluster 6 with Supercell 62 excluded. Examined in isolation, Supercell 62 displayed a coherent transcriptional programme characteristic of conventional dendritic cells (cDC2). It showed strong up-regulation of classical DC markers (*CD1C, CD1E, FCER1A, CLEC10A*) together with MHC class II genes (*HLA-DPA1, HLA-DQA1, HLA-DPB1, CD74*). Strikingly, these genes were not significantly enriched when Cluster 6 was considered as a whole, indicating that their signal was largely masked at the cluster level. Additional genes—including *MS4A6A, IRF4, NDRG2*, and *LINC00926* —were likewise elevated only in Supercell 62, further substantiating its distinct identity and antigen-presenting potential. When Supercell 62 was excluded from Cluster 6, all dendritic-cell markers lost statistical significance, and the residual cluster reverted to a CD8^+^ T-cell–like or otherwise heterogeneous phenotype. This analysis demonstrates that the apparent cDC2-like programme within Cluster 6 was driven almost entirely by Supercell 62.

To further delineate the biological processes that distinguish Supercell 62, we performed gene-set enrichment analysis (GSEA) across four pathway resources—Gene Ontology (GO), Reactome, MSigDB, and KEGG (Supplementary Table 3).

#### GO

Gene Ontology enrichment revealed significant over-representation of vesicular-membrane terms involved in antigen trafficking and presentation, including *ER-to-Golgi transport vesicle membrane* (NES = 2.05, *q* = 0.0106), *clathrin-coated endocytic vesicle membrane* (NES = 2.04), and *transport vesicle membrane* (NES = 2.03). All three terms were driven by a common core of MHC-II genes—*HLA-DRA, HLA-DPA1, HLA-DQA1, CD74*, and related loci.

#### Reactome

Antigen-presentation pathways were likewise prominent, with *MHC class II antigen presentation* (NES = 2.02, *q* = 0.0179) leading the list; components of *TCR signalling* and *IFN-γ signalling* were also enriched, again centred on MHC-II genes, underscoring the antigen-presenting capacity of Supercell 62.

#### MSigDB

Convergent evidence was obtained from curated gene sets, including *Hay Bone Marrow Dendritic Cell* (NES = 2.02, *q* = 1.9 × 10^−8^) and *Transport Vesicle Membrane* (NES = 2.08), both supporting a dendritic-cell phenotype.

#### KEGG

Pathways related to antigen-driven immunity and immunopathology were highlighted, such as *Intestinal immune network for IgA production* (NES = 2.09), *Systemic lupus erythematosus*, and *Graft-versus-host disease*; each was enriched by overlapping HLA-class II genes, further consistent with dendritic-cell–mediated immune functions.

To complement the GSEA findings, we constructed an integrated multi-panel figure (Fig. 5) summarising pathway enrichment across four knowledge bases. The cross-database visualisations corroborate our findings and substantiate the interpretation of Supercell 62 as a mature, functionally autonomous cDC2 population. Recent work has demonstrated that the cDC2 is functionally heterogeneous, comprising molecularly and ontogenetically distinct sub-lineages [44, 45]. This diversity implies that only high-resolution approaches such as supracellular modelling can mitigates the signal “dilution” observed at the cluster level, thereby disentangle rare or specialised cell populations.

**Fig 5.**
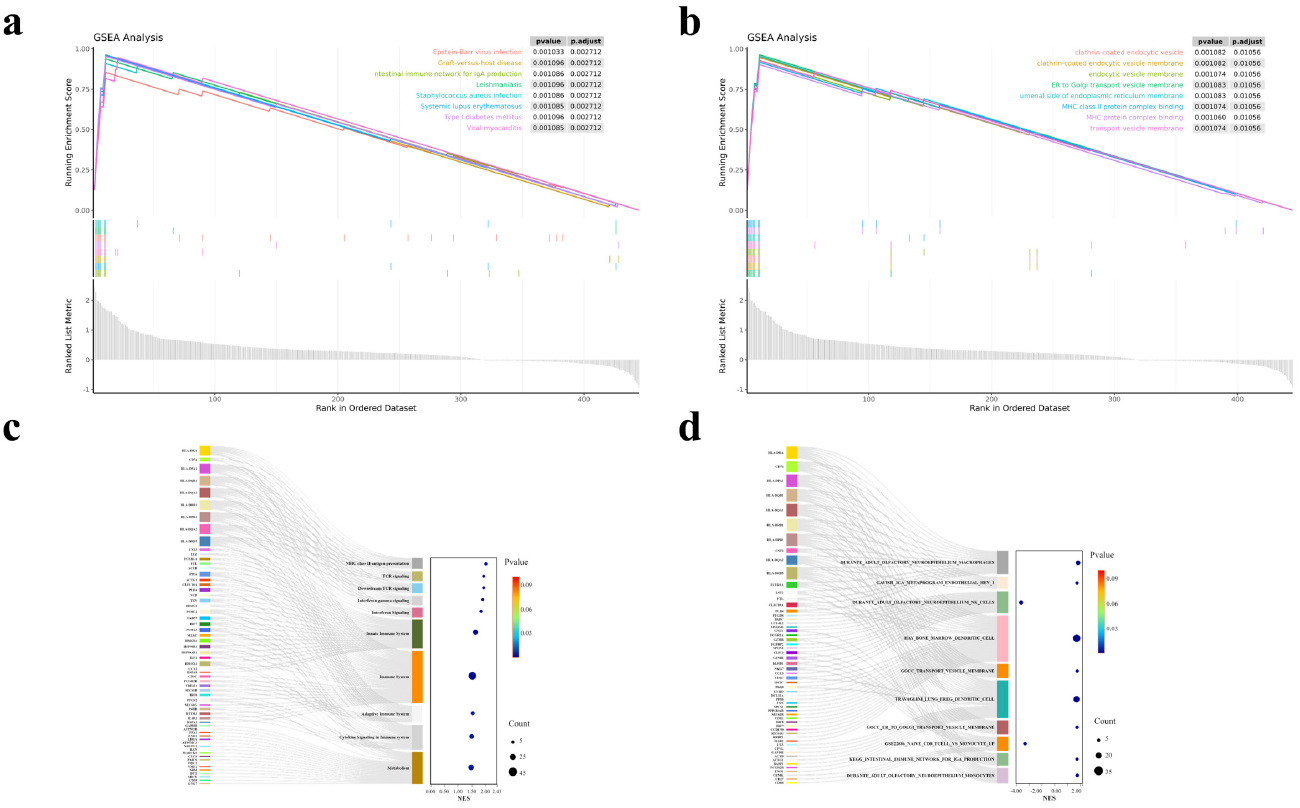
Pathway enrichment analysis for Supercell 62 across four databases. Each panel displays the ranked gene set enrichment scores (GSEA) or gene-pathway mapping (Sankey plots), emphasizing coherent activation of MHC-II mediated antigen presentation, dendritic cell programs, vesicle trafficking, and immune signaling pathways. (a) KEGG pathway enrichment. It pointed to specialised immune functions, highlighting pathways such as *intestinal immune network for IgA production* and *graft versus host disease*, both of which depend on antigen-presenting activity.(b) GO component enrichment. Membrane compartments essential to antigen processing and trafficking—such as *clathrin-coated vesicles* and *ER-to-Golgi transport vesicles* —were significantly enriched. (c) Reactome pathway enrichment. It accentuated *MHC class II antigen presentation* together with downstream *TCR* and *IFN-γ* signalling cascades. (d) MSigDB enrichment. It showed strong concordance with curated dendritic-cell programmes, notably *Hay Bone Marrow Dendritic Cell*, as well as with vesicle-associated signatures.

Another instance of a rare cell population revealed by supercell modelling is exemplified by Supercell 363, the largest mesoscopic unit embedded within Cluster 5, which is annotated as a B-cell population (Supplementary Table 4). A comparative analysis—Supercell 363 and Cluster 5 with Supercell 363 excluded—showed that the NK-cell–associated surface proteins CD56 and TIGIT were significantly up-regulated only in Supercell 363. These signals were diluted in the full cluster and were lost once Supercell 363 was excluded, indicating that NK-related expression is confined to this supracellular unit and masked at the bulk-cluster level. The coexistence of both NK and B cell markers within a single supercell suggests a rare composite phenotype, which were previously reported as rare natural killer-like B cells (NKB) [46].

Together, these examples illustrate that by aggregating transcriptionally and proteomically coherent cells, supracellular modelling can reveal atypical populations, underscoring its ability to resolve rare or functionally specialised cell populations that would otherwise remain concealed.

### t-SNE projections expose rare populations captured by supercells

To illustrate the utility of supracellular abstraction, we mapped Supercell 62 and Supercell 363 onto the t-SNE projection of the joint single-cell multi-omics manifold.

As illustrated in Fig. 6a, Supercell 62 (black triangles) aligns almost exclusively with a sparsely distributed dendritic-cell cohort that remains invisible in any single-cell clustering partition. Conventional clustering thus under-samples or fragments this rare population, whereas supracellular abstraction consolidates it into a expression-coherent unit, exposing its presence at the mesoscopic scale and demonstrating the utility of supercells for recovering noise-masked minority cell types.

**Fig 6.**
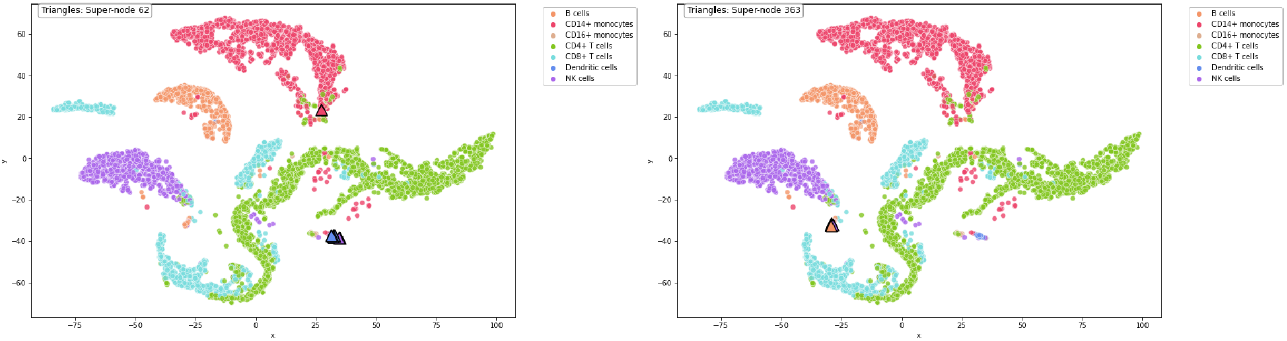
Visualization of supercell regions in t-SNE embedding. Both supercells reside in spatially isolated areas, illustrating the ability of supercell-level resolution to uncover rare and intermediate biological states. (a) Supercell 62 highlights a rare dendritic cell group not detectable by conventional clustering. (b) Supercell 363 captures a group of B cells with two embedded NK cells, representing a rare composite population.

As shown in Fig. 6b, Supercell 363 occupies a distinct t-SNE “island” composed chiefly of B cells together with two NK cells. Its clear separation from the main B-cell cluster implies a rare composite population corresponding to the previously described NK-like B cells ([46]). By consolidating these boundary-spanning cells into a single module, the supracellular framework remains sensitive to biologically meaningful heterogeneity that would otherwise be lost at the margins of canonical lineages.

In both cases, the supercells occupy spatially isolated regions that are clearly distinct from the major cellular aggregates. This exemplifies a key advantage of supracellular modelling: its capacity to capture intermediate-scale, structurally specific units that lie between single-cell resolution and cluster granularity. Such units provide a powerful abstraction for uncovering hidden biological states within large-scale single-cell datasets.

### scHG achieves state-of-the-art multi-omics clustering performance across six benchmarks

We conducted comprehensive evaluations on six public datasets (see Section), benchmarking scHG against six state-of-the-art approaches (see Section) using the **ARI** and **NMI** metrics. Quantitative comparisons are presented in Tables 2 and 3.

**Table 2.** Performance comparison of different methods in **ARI** on the considered datasets.

| Datasets<br>Methods | PBMC10× | PBMC_Inhouse | Sim | Bmcite | mESC | PBMC_Cao |
| --- | --- | --- | --- | --- | --- | --- |
| GBS | 0.6750 | 0.4642 | **0.9881** | — | 0.6881 | 0.4231 |
| scAI | 0.7208 | 0.6598 | 0.9239 | — | **0.8629** | **0.4642** |
| SMSC | 0.6981 | 0.6320 | **0.9881** | — | 0.7329 | 0.3756 |
| CGD | **0.8078** | **0.6914** | 0.6901 | — | 0.6042 | 0.4512 |
| scMNMF | 0.7716 | 0.5950 | 0.7278 | — | 0.8226 | 0.4153 |
| GSTRPCA | 0.5621 | 0.6295 | 0.8445 | — | — | — |
| scHG | **0.8774** | **0.7029** | **0.9646** | **0.6732** | **0.9373** | **0.5299** |

**Table 3.** Performance comparison of different methods in **NMI** on the considered datasets.

| Datasets<br>Methods | PBMC10× | PBMC_Inhouse | Sim | Bmcite | mESC | PBMC_Cao |
| --- | --- | --- | --- | --- | --- | --- |
| GBS | 0.7100 | 0.4443 | **0.9832** | — | 0.6440 | 0.4966 |
| scAI | 0.6617 | 0.7302 | 0.9425 | — | **0.8014** | 0.4749 |
| SMSC | 0.6954 | 0.6930 | **0.9832** | — | 0.7112 | 0.5024 |
| CGD | 0.7458 | **0.7989** | 0.6883 | — | 0.5873 | **0.5283** |
| scMNMF | **0.7505** | 0.6468 | 0.7544 | — | 0.6564 | NaN |
| GSTRPCA | 0.4685 | 0.6451 | 0.8614 | — | — | — |
| scHG | **0.8230** | **0.7992** | **0.9598** | **0.6917** | **0.8589** | **0.5972** |

scHG achieves superior performance on most benchmark datasets (PBMC10×, PBMC Inhouse, Bmcite, mESC, PBMC Cao), demonstrating leading **ARI** and **NMI** values. On sim dataset, although ranking third in both metrics, scHG attains competitive third-place rankings with both **ARI** and **NMI** exceeding 95%, closely approaching optimal clustering performance. Notably, the strong alignment between **ARI** and **NMI** rankings confirms our improvements originate from intrinsic high-order structural preservation rather than metric-specific optimization.

To evaluate embedding quality, we project high-dimensional omics data from all six benchmark datasets into two-dimensional space using t-SNE. Figs. 7-11 display visualizations for five datasets (except dataset bmcite to which all the comparision methods are inapplicable).^3^

**Fig 7.**
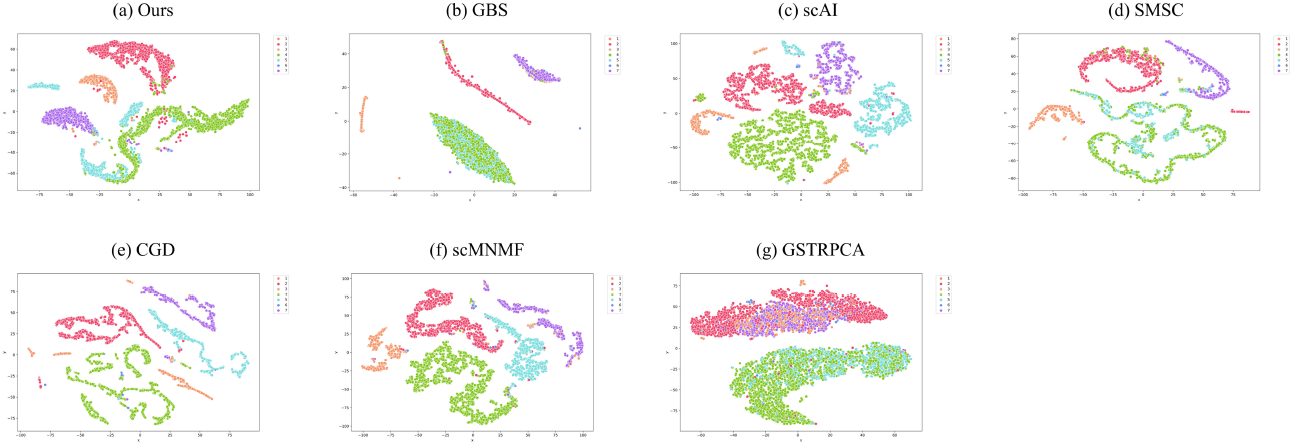
Comparative t-SNE Visualization of Latent Representations on PBMC10× Dataset: (a) scHG, (b) GBS, (c) scAI, (d) SMSC, (e) CGD, (f) scMNMF, (g) GSTRPCA.

**Fig 8.**
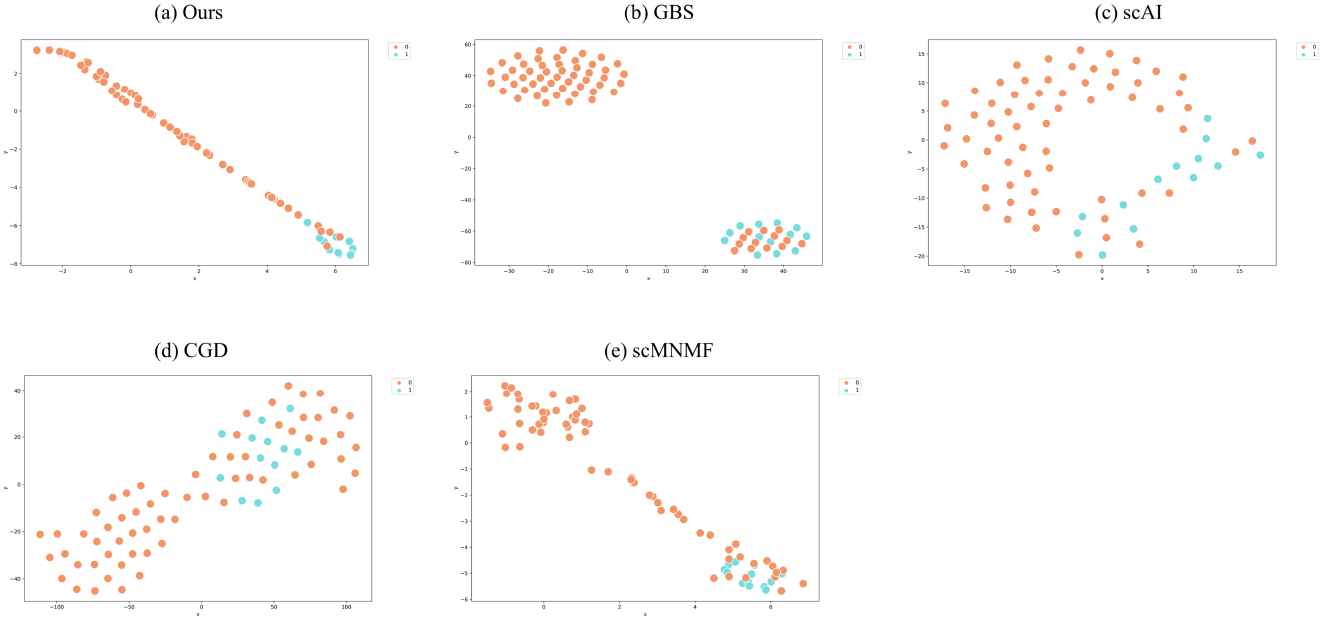
Comparative t-SNE Visualization of Latent Representations on mESC Dataset: (a) scHG, (b) GBS, (c) scAI, (d) CGD, (e) scMNMF.

**Fig 9.**
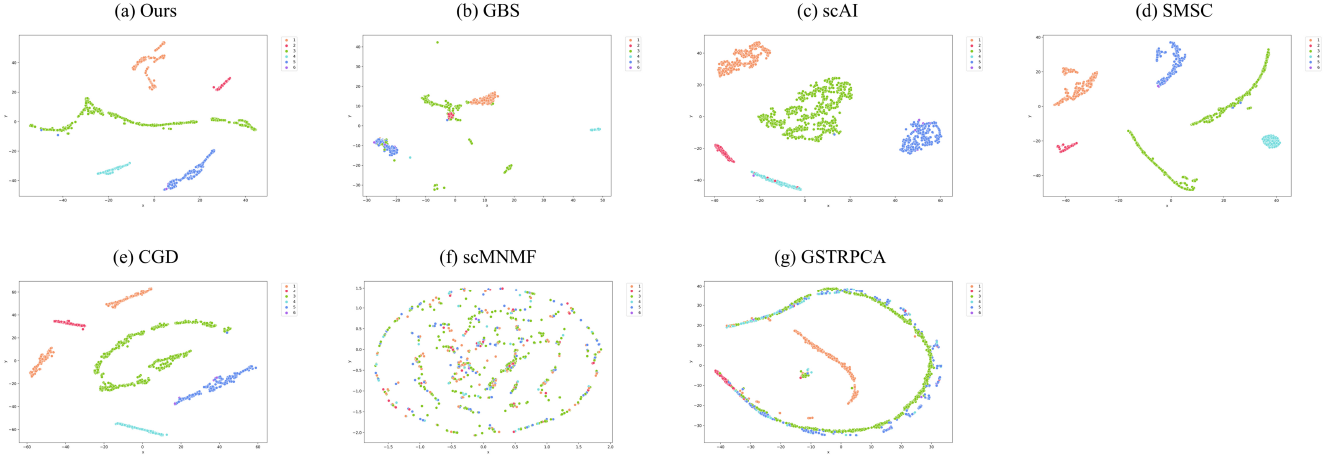
Comparative t-SNE Visualization of Latent Representations on pbmc inhouse Dataset: (a) scHG, (b) GBS, (c) scAI, (d) CGD, (e) scMNMF.

**Fig 10.**
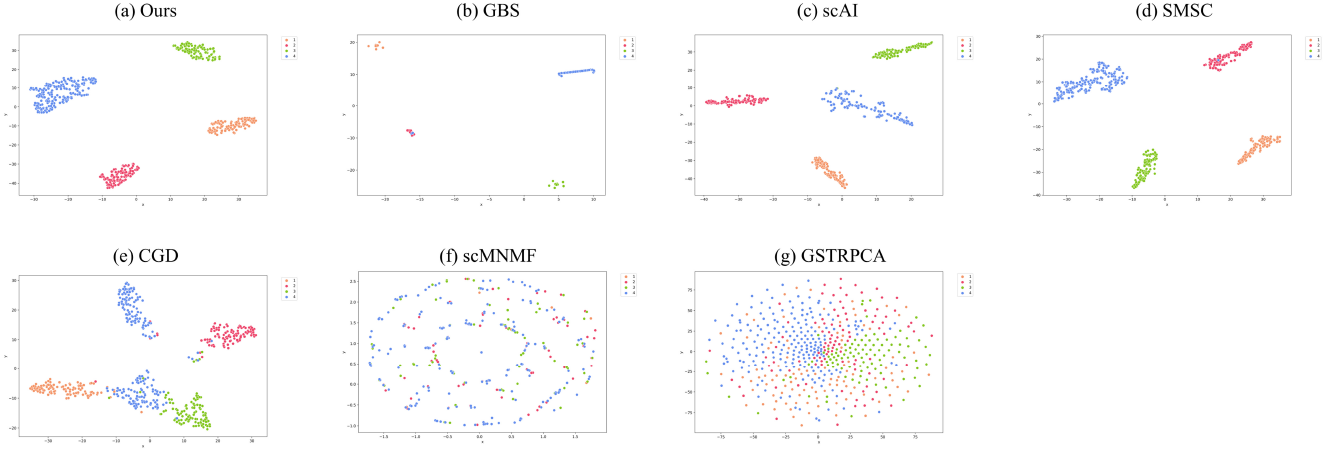
Comparative t-SNE Visualization of Latent Representations on sim Dataset: (a) scHG, (b) GBS, (c) scAI, (d) CGD, (e) scMNMF.

**Fig 11.**
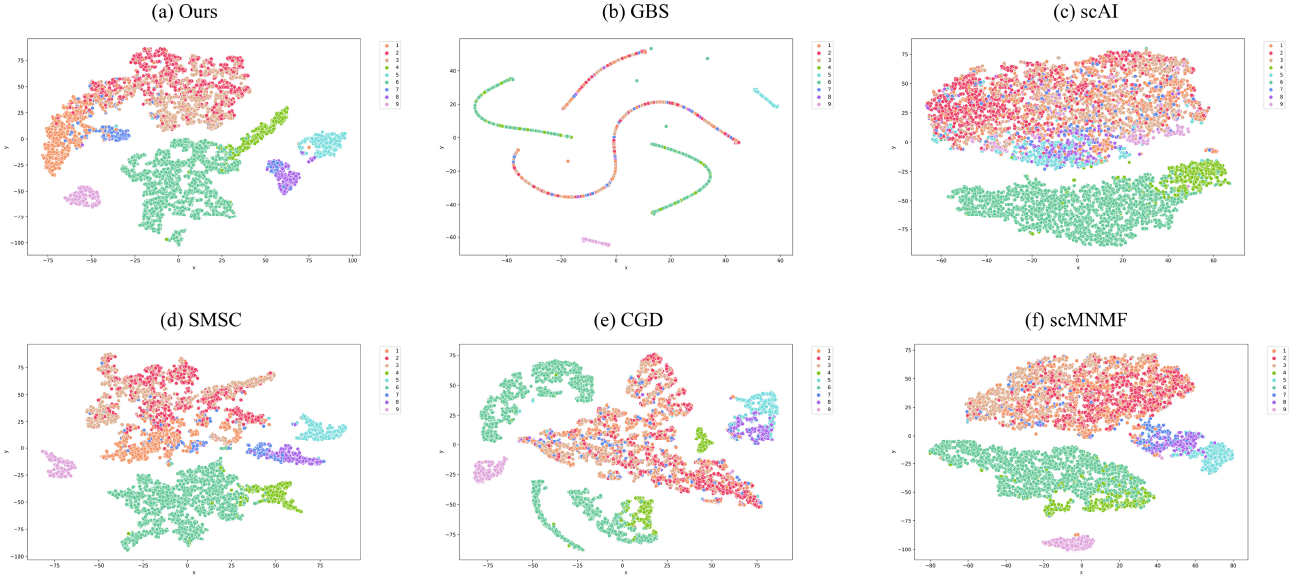
Comparative t-SNE Visualization of Latent Representations on pbmc Cao Dataset: (a) scHG, (b) GBS, (c) scAI, (d) CGD, (e) scMNMF.

Take the PBMC10× dataset as an example. scHG achieves tight, well-separated clusters that corresponding to distinct cell types, outperforming baselines in structural preservation: (c) scAI (e) CGD (f) scMNMF exhibit high intra-cluster dispersion. (b) GBS (g) GSTRPCA exhibit significant inter-cluster overlap. (d) SMSC exhibit both intra-cluster fragmentation and inter-cluster overlap.

Across all five datasets, scHG consistently demonstrates compact intra-cluster structure and well-separated inter-cluster boundaries in the visualization results. By aggregating cells with consistent multi-hop neighborhoods, supercells promote intra-cluster compactness. Simultaneously, pruning step eliminates weak or spurious intra-supercell links, thereby enhancing inter-cluster separability by minimizing false connections between unrelated regions.

### High-order supercell strategy achieves state-leading runtime efficiency on large-scale multi-omics datasets

We benchmarked computational efficiency across six comparative methods and our approach on public datasets (Table 4 & Fig 12).

**Table 4.** Performance comparison of different methods in time(s) on the considered datasets.

| Methods \ Datasets | PBMC10× | PBMC_Inhouse | Sim | Bmcite | mESC | PBMC_Cao |
| --- | --- | --- | --- | --- | --- | --- |
| GBS | 266.324 | 6.546 | <u>1.036</u> | — | **0.309** | 562.208 |
| scAI | 14771.801 | 271.596 | 47.234 | — | 1.167 | 26060.514 |
| SMSC | 90.112 | <u>3.823</u> | **0.581** | — | <u>0.362</u> | 95.281 |
| CGD | 1154.190 | 909.180 | 8.617 | — | 0.442 | 3191.165 |
| scMNMF | <u>31.957</u> | 40.033 | 3.691 | — | 0.905 | <u>53.151</u> |
| GSTRPCA | 1956.157 | 2000.284 | 392.497 | — | — | — |
| scHG | **13.781** | **3.064** | 1.112 | **1104.834** | 0.518 | **32.507** |

<sup>3</sup>GSTRPCA is inapplicable to datasets mESC and PBMC\_Cao, and SMSC yields a 1-dimensional latent space for this dataset; consequently, both methods lack t-SNE visualizations in the dataset mentioned above.

**Fig 12.**
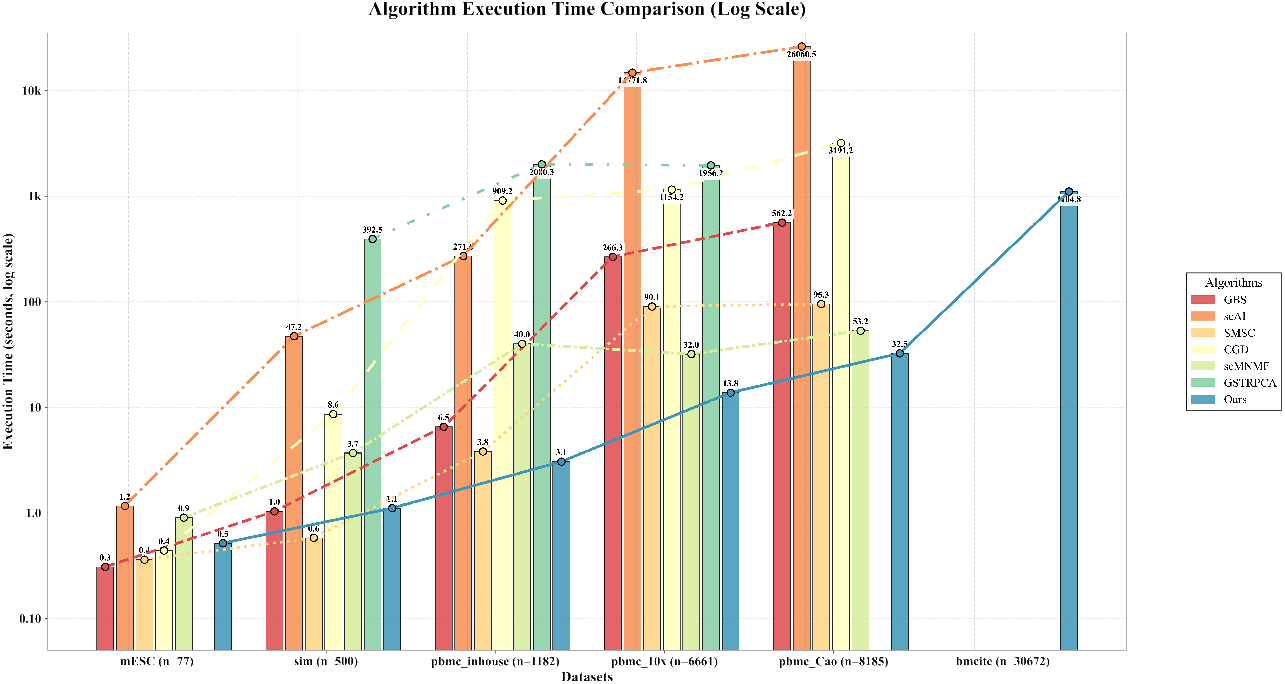
Execution time comparison (Log Scale) of different methods on the considered datasets.

Across smaller-scale datasets (mESC, n=77; Sim, n=500), scHG demonstrated competitive efficiency, ranking as the fourth and third fastest solution respectively. Conversely, for larger datasets including PBMC Inhouse (n=1182), PBMC10× (n=6661), PBMC Cao (n=8185), and Bmcite (n=30672), it achieved superior execution speeds. Figure 12 quantifiably confirms the progressive enhancement of scHG’s runtime advantage with increasing sample sizes.

This performance aligns with computational complexity analysis: our *O*(|*V*| ^2^) algorithm (see the time complexity analysis in Section) avoids more complex computations required by comparative methods, particularly beneficial for large-scale datasets. The efficiency gains originate from our supercell construction and two-stage clustering framework, which reduces computational load compared to primitive cell processing.

### High-order refinement enhances supercell purity by correcting second-order misclassifications

To illustrate the benefit of incorporating high-order information, we analysed the PBMC10× dataset (6661 cells, Supplementary Table 5). Using only second-order co-occurrence, the fused cross-omic graph partitions the data into 4802 supercells.

Among these, seven supercells (IDs 62, 63, 363, 453, 1271, 1852, 1917) contain at least three cells and span two or more ground-truth labels. After augmenting the graph with higher-order neighborhoods and degree-centrality pruning, many misassigned cells are removed. We quantify this effect with **Recall**_*sc*_:

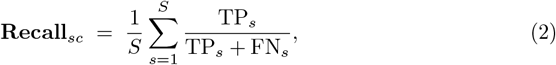

where TP_*s*_ and FN_*s*_ counts, respectively, denote the mislabelled cells successfully pruned and those that remain in supercell *s*. The confusion matrix in Table 5 yields a **Recall**_*sc*_ of 75.97 %, indicating that three-quarters of the second-order misclassifications are corrected by the high-order refinement.

**Table 5.**
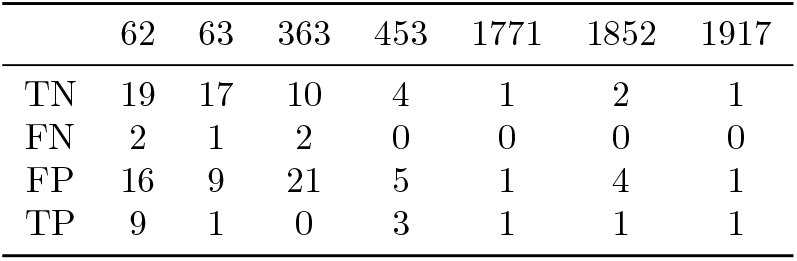
Cell–level confusion statistics for the seven supercells (IDs 62, 63, 363, 453, 1771, 1852, 1917) in the PBMC10× dataset before and after high-order refinement.

|  | 62 | 63 | 363 | 453 | 1771 | 1852 | 1917 |
| --- | --- | --- | --- | --- | --- | --- | --- |
| TN | 19 | 17 | 10 | 4 | 1 | 2 | 1 |
| FN | 2 | 1 | 2 | 0 | 0 | 0 | 0 |
| FP | 16 | 9 | 21 | 5 | 1 | 4 | 1 |
| TP | 9 | 1 | 0 | 3 | 1 | 1 | 1 |

Fig. 13 visualises Supercell 62. Of eleven cells initially mis-grouped, nine are eliminated by the high-order step; these cells have markedly lower degree centrality than retained members, confirming that centrality is an effective proxy for confidence. Overall, the high-order strategy substantially increases supercell purity and underscores its value for dissecting single-cell multi-omics heterogeneity.

**Fig 13.**
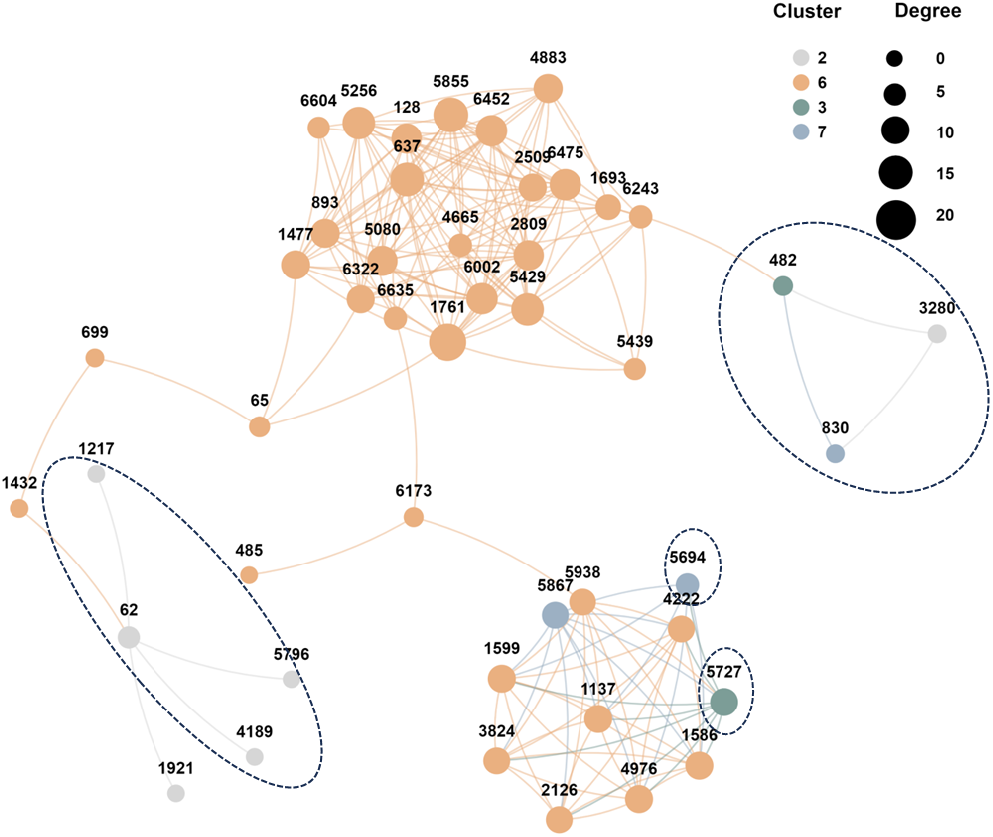
Connectivity graph of Supercell 62 obtained from the second-order partition before high-order pruning. Node size is proportional to degree centrality; colours indicate ground-truth clusters. Cells outlined by dashed ellipses are the low-centrality outliers that are removed by the high-order refinement, illustrating how the pruning step improves supercell purity by eliminating mis-assigned cells. (This plot were generated using the CNSknowall platform (https://cnsknowall.com), a comprehensive web service for data analysis and visualization.)

## Discussion

### Ablation analysis

To validate the effectiveness of the proposed angle-aware metric and probabilistic pruning modules, we performed detailed ablation studies across all six datasets (referenced in Section). Our experimental protocol establishes a baseline model employing Euclidean distance without probabilistic pruning, maintaining identical hyperparameters and solution methodologies to ensure comparative fairness.

Quantitative performance improvements are evaluated across eight metrics in Table 6: Accuracy (**ACC**; [47]), Normalized Mutual Information (**NMI**; [48, 49]), **Purity** ([50]), **F1-score** ([51]), **Precision** ([51]), **Recall** ([51]), Rand Index (**RI**; [52]), and Adjusted Rand Index (**ARI**; [53]). In particular, we present the comparison of **ARI** and **NMI** through Fig. 14.

**Table 6.**
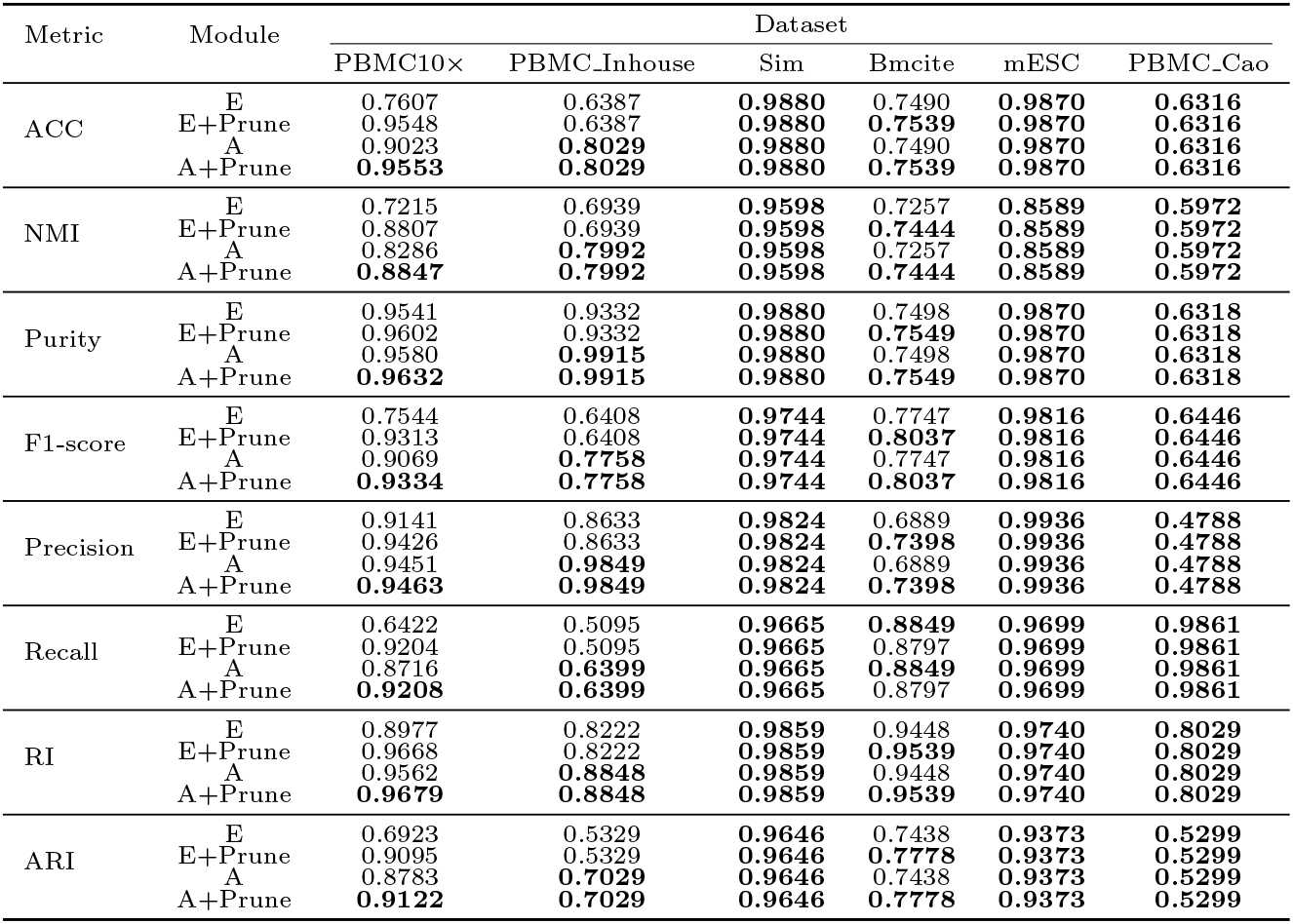
Ablation results of angle-aware metric and probabilistic pruning modules in six datasets across eight clustering evaluation metrics; E stands for Euclidean metric and A for angle-aware metric.

**Fig 14.**
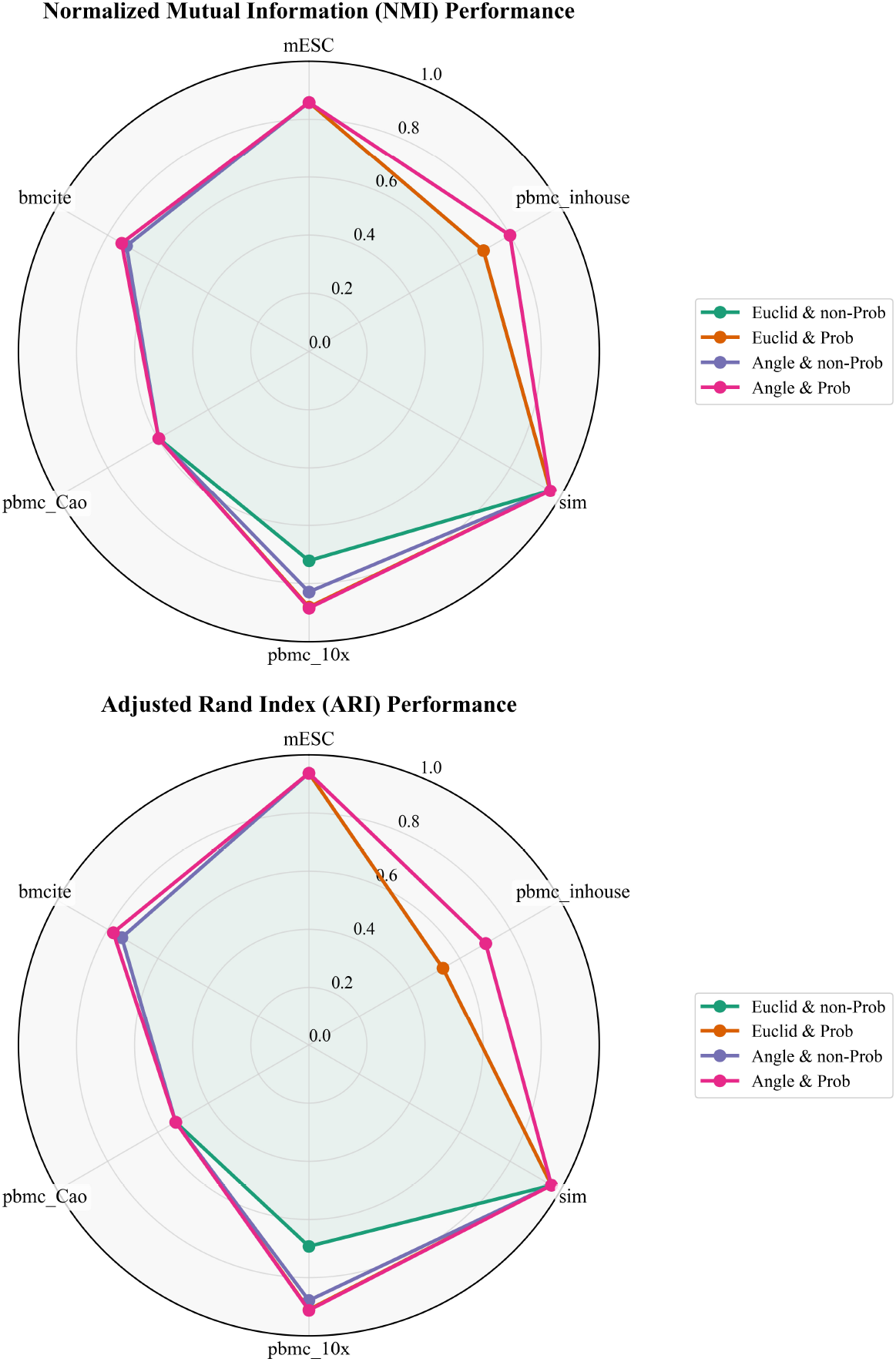
Ablation experiment results (**NMI & ARI**) on the considered datasets.

For PBMC10×, PBMC Inhouse, and Bmcite datasets, models incorporating angle-aware metrics and probabilistic pruning demonstrated substantial performance gains relative to Euclidean-distance counterparts without pruning: **NMI** improvements of 16.32%, 10.53%, and 1.87%; and **ARI** improvements of 21.99%, 17.00%, and 3.40% respectively. On the remaining three datasets, these components maintained performance without degradation. Collectively, our ablation results confirm that each proposed component contributes non-negligibly to performance enhancement, supporting the overall efficacy of scHG design.

### Performance on different cluster numbers

We conducted systematic evaluations of clustering sensitivity across benchmark datasets by measuring **ARI**/**NMI** variation with cluster number (Figs. 15-16). On four datasets (PBMC Inhouse, Sim, mESC, PBMC Cao), scHG achieves exact alignment between algorithm-optimized (green) and ground-truth (yellow) cluster numbers, with identical **ARI**/**NMI** rankings. For the remaining two datasets (PBMC10×, Bmcite), while minor discrepancies existed in cluster number estimation, our model achieved indistinguishable performance compared to ground-truth configurations.

**Fig 15.**
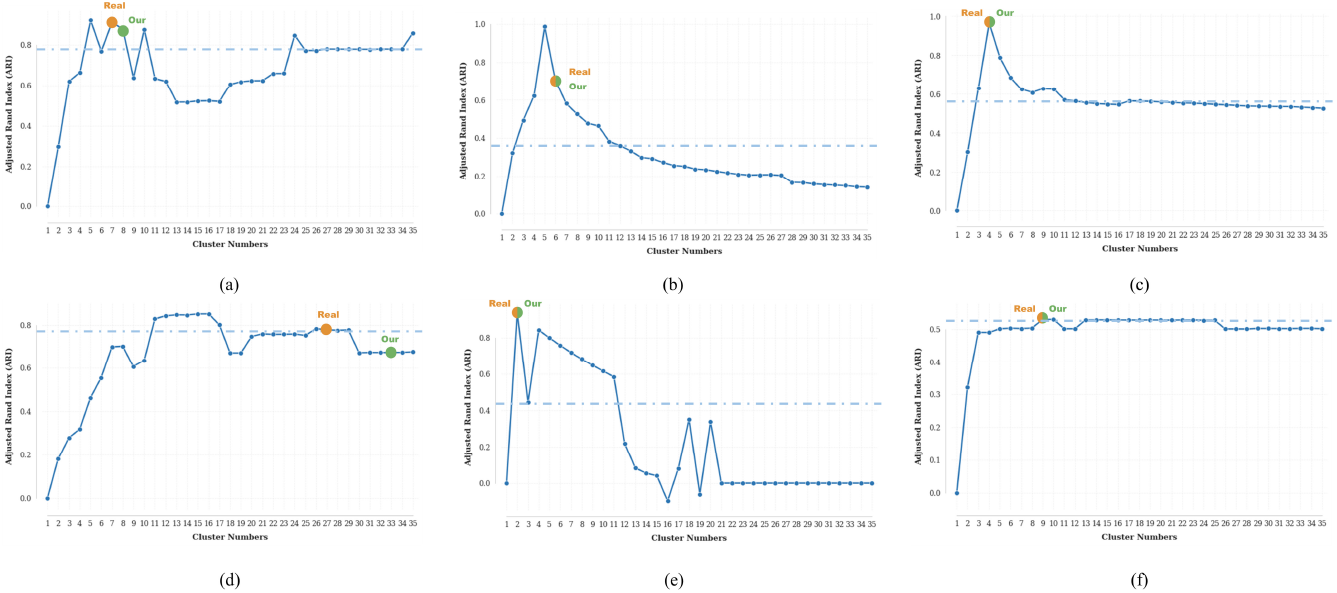
**ARI** variation across six benchmark datasets: (a) PBMC10×, (b) PBMC Inhouse, (c) Sim, (d) Bmcite, (e) mESC, (f) PBMC Cao. Dots indicate: yellow (ground-truth cluster numbers), green (algorithm-derived optimal cluster numbers). Dashed indicate: blue (top-10 ARI threshold).

**Fig 16.**
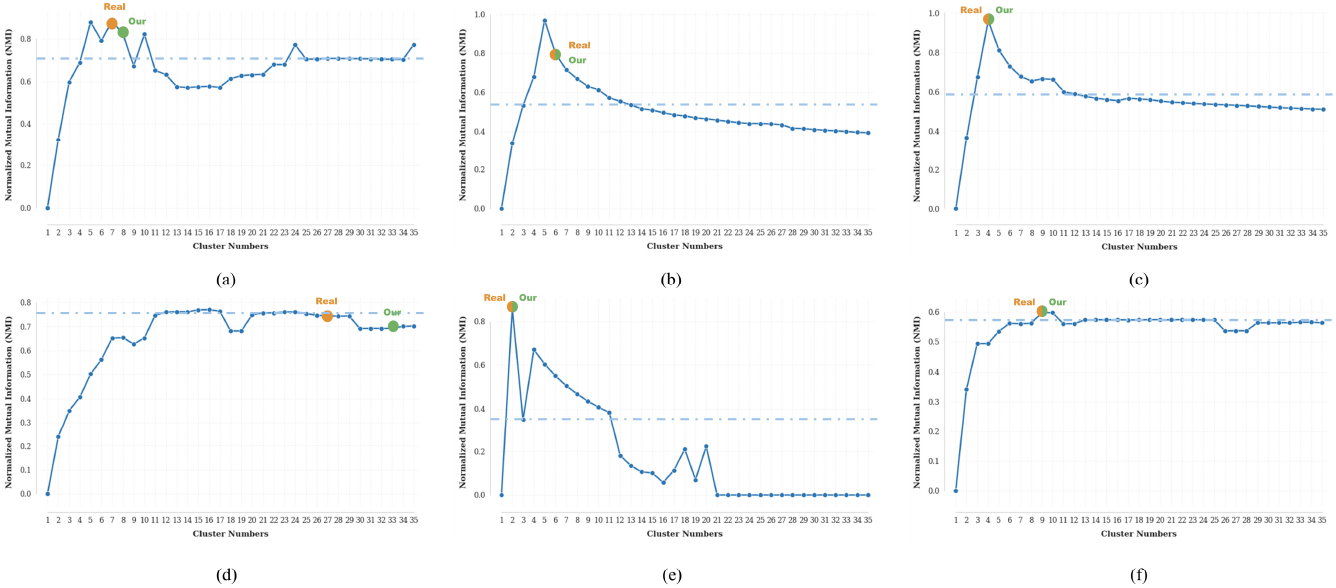
**NMI** variation across six benchmark datasets: (a) PBMC10×, (b) PBMC Inhouse, (c) Sim, (d) Bmcite, (e) mESC, (f) PBMC Cao. Dots indicate: yellow (ground-truth cluster numbers), green (algorithm-derived optimal cluster numbers). Dashed indicate: blue (top-10 NMI threshold).

Across half of the benchmark datasets (PBMC10×, Bmcite, PBMC Cao), the top-10 thresholds achieve 84.6–99.6% of the maximum **ARI** and 80.0–98.3% of the maximum **NMI**, indicating high robustness to cluster number selection.

For the remaining datasets (PBMC Inhouse, Sim, mESC), the top-10 thresholds achieve 36.6–58.5% of the maximum **ARI** and 40.6–61.2% of the maximum **NMI**, while maintaining 100% accuracy in cluster number estimation (Fig. 15–16), underscoring the method’s capability to precisely determine cluster numbers across sensitivity ranges.

## conclusion

We present scHG, a high-order supercell framework that achieves both scalability and subtype-aware resolution in multi-omics integration. By abstracting expression-coherent cells into supercells, scHG applies an angle-aware metric and a modality-weighted block coordinate descent optimization to cluster supercells efficiently.

scHG processes 30000 cells in under 20 minutes, consistently delivering state-of-the-art ARI and NMI across five real-world datasets. Without relying on prior cluster labels, it retrieves clinically relevant B-cell targets and resolves heterogeneous subpopulations within reference-defined T-cell clusters. The supercell perspective further exposes rare dendritic-cell clusters and NK-like B-cell intermediates that conventional pipelines fail to detect. Ablation studies demonstrate that the high-order graph model—integrating second-order co-occurrence neighbors, angle-aware metrics, and stochastic pruning—boosts performance by an average of 14% across eight clustering metrics relative to second-order graph baselines, while enabling adaptive determination of cluster numbers without prior assumptions.

Together, these findings underscore the power of the supercell paradigm, grounded in high-order graph representations, to uncover rare immune states and decode tissue complexity. Its inherently graph-based architecture provides a natural avenue for incorporating spatial context and additional omics layers, offering a forward-looking analytical framework with strong potential for deeper biological insight and clinically actionable discovery.

## Materials and methods

Consider a collection of *V* multi-omics datasets 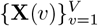, where each matrix 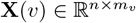 represents a distinct omics data. Here *n* denotes the number of cells, and *m*_*v*_ the number of features for the *v*-th omics data **X**(*v*). Let **x**^*i*^(*v*) denote the *i*-th row vector (*i*-th cell) and **x**_*j*_(*v*) the *j*-th column vector (*j*-th feature) of **X**(*v*), corresponding to biological features such as gene expression (RNA-seq), surface protein abundance (antibody-derived tags (ADT)), or chromatin accessibility (ATAC-seq).

### Angle-aware metric

In traditional omics clustering analyses, similarity between cells is frequently quantified using Euclidean distance, whereby smaller distances indicate greater similarity. In contrast, Eq. (3) defines an alternative angle-aware metric: the Pearson correlation coefficient (PCC) between cells.

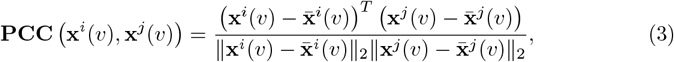

Where **x**^*i*^(*v*) and **x**^*j*^(*v*) are any two cells in the *v*-th omics, respectively.

Building on the angle-aware metric introduced earlier, we can distinguish cell pairs that are geographically proximate (small Euclidean distance) but lack strong linear correlation. As illustrated in Fig. 17, *d*_1_ *> d*_2_ indicates greater Euclidean separation between cells *i* and *j* than between *i* and *k*, implying lower Euclidean-based similarity. Consequently, these cells (*i* and *k*) would typically be grouped together under traditional clustering. However, the angular metric reveals that *θ*_1_ *< θ*_2_, leading to **PCC** (*x*^*i*^, **x**^*j*^) *>* **PCC** (**x**^*i*^, **x**^*k*^). This demonstrates significantly higher linear-correlation similarity between cells *i* and *j* than between *i* and *k*, which is more reasonable.

**Fig 17.**
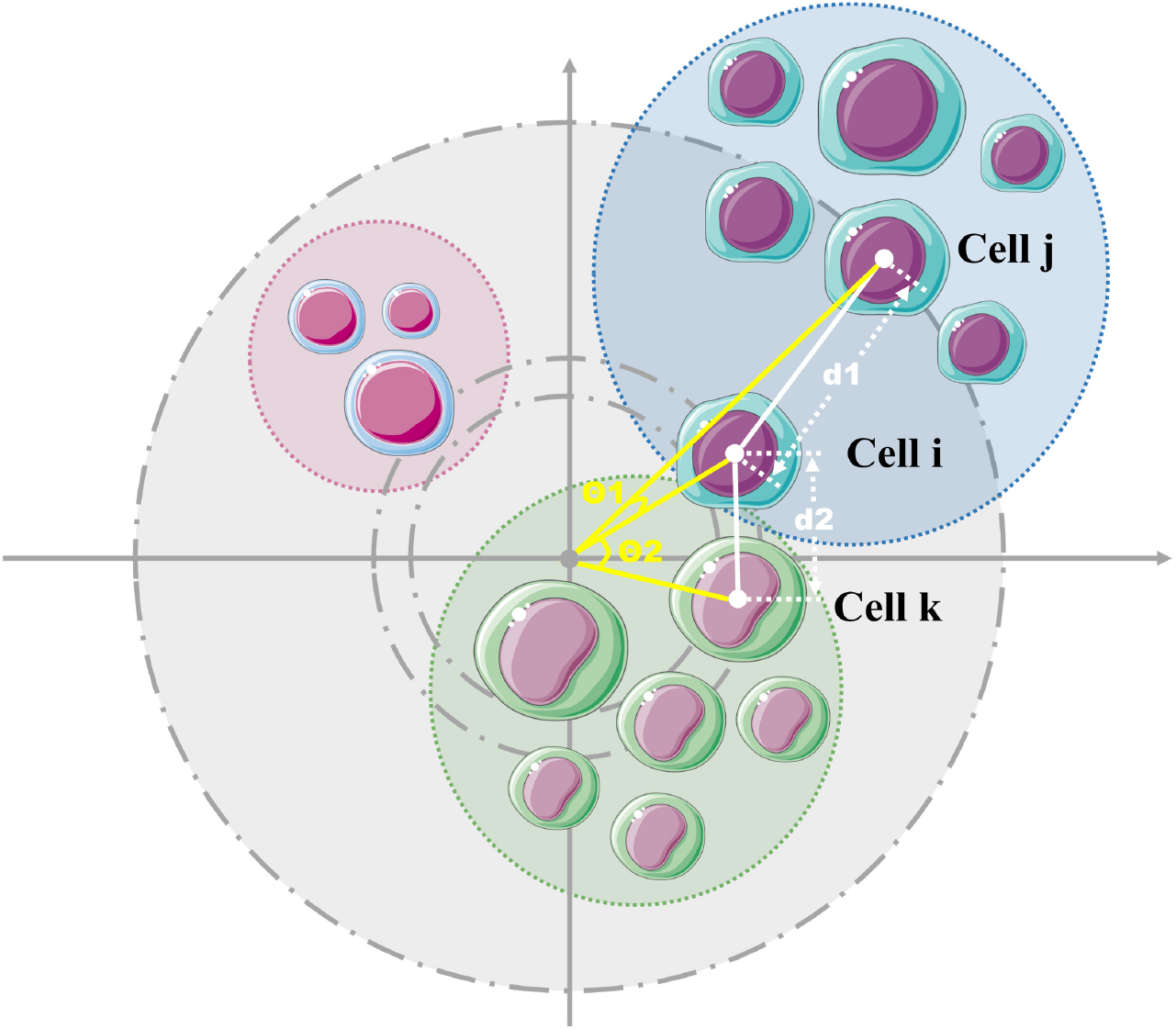
Angle-aware metric versus Euclid metric. According to the picture, *d*_1_ *> d*_2_ indicates greater Euclidean separation between cells *i* and *j* than between *i* and *k*, implying lower Euclidean-based similarity. Consequently, these cells (*i* and *k*) would typically be grouped together under traditional clustering. However, the angular metric reveals that *θ*_1_ *< θ*_2_, leading to **PCC (x**^*i*^, **x**^*j*^ *)>* **PCC (x**^*i*^, **x**^*k*^ ). This demonstrates significantly higher linear-correlation similarity between cells *i* and *j* than between *i* and *k*, which is more reasonable.

Given these comparative advantages of the angle-aware metric over Euclidean distance, our algorithm employs angle-aware metric as the primary similarity measure.

### High order neighbor-aware coarsening graph clustering framework

The cell similarity matrix **A**(*v*) = (*a*_*ij*_)_*n*×*n*_ for the *v*-th omic is constructed using the *γ*-proximity linear similarity metric. Specifically, we first compute the Pearson correlation matrix **PCC**(*v*) of matrix **X**(*v*) by:

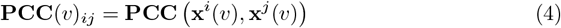

Then for each cell *i*, we extract its *γ*+1 nearest neighbors (excluding self) from **PCC**(*v*) to form the truncated ordered distance matrix 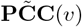, yielding the similarity coefficients:

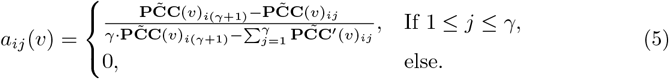

Given *C* cell clusters, let 𝒞_*p*_ denote the *p*-th cluster. The inter-cluster similarity between 𝒞_*p*_ and its complement 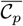 is defined as:

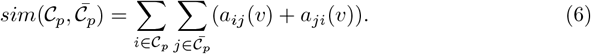

The total inter-cluster similarities across all clusters and their complements is quantified as:

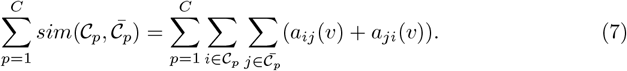

To account for omics-specific contributions, we introduce modality weights 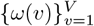, yielding the weighted multi-omics similarity:

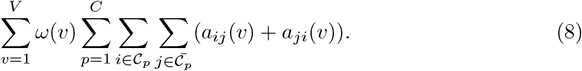

To seek maximally separable partitions, the total multi-omics similarity should be minimized through the following constrained optimization:

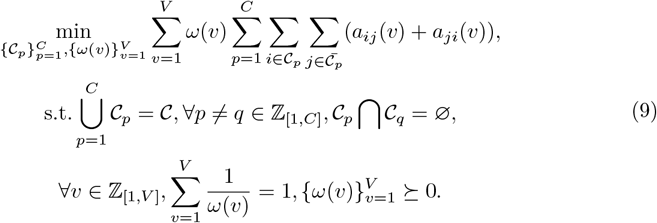

To capture the complex similarity relationships among cells, we introduce second-order information. For the *v*-th omic, let 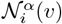 denote the *α*-nearest neighbors of cell *i*, and **N**(*v*) ∈ {0, 1} ^*n*×*n*^ represent the second-order co-occurrence matrix where elements are determined by:

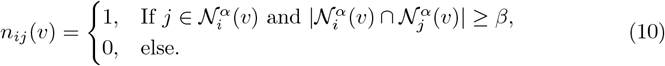

An example of second-order co-occurrence neighbors is visualized in Fig. 18(b).

**Fig 18.**
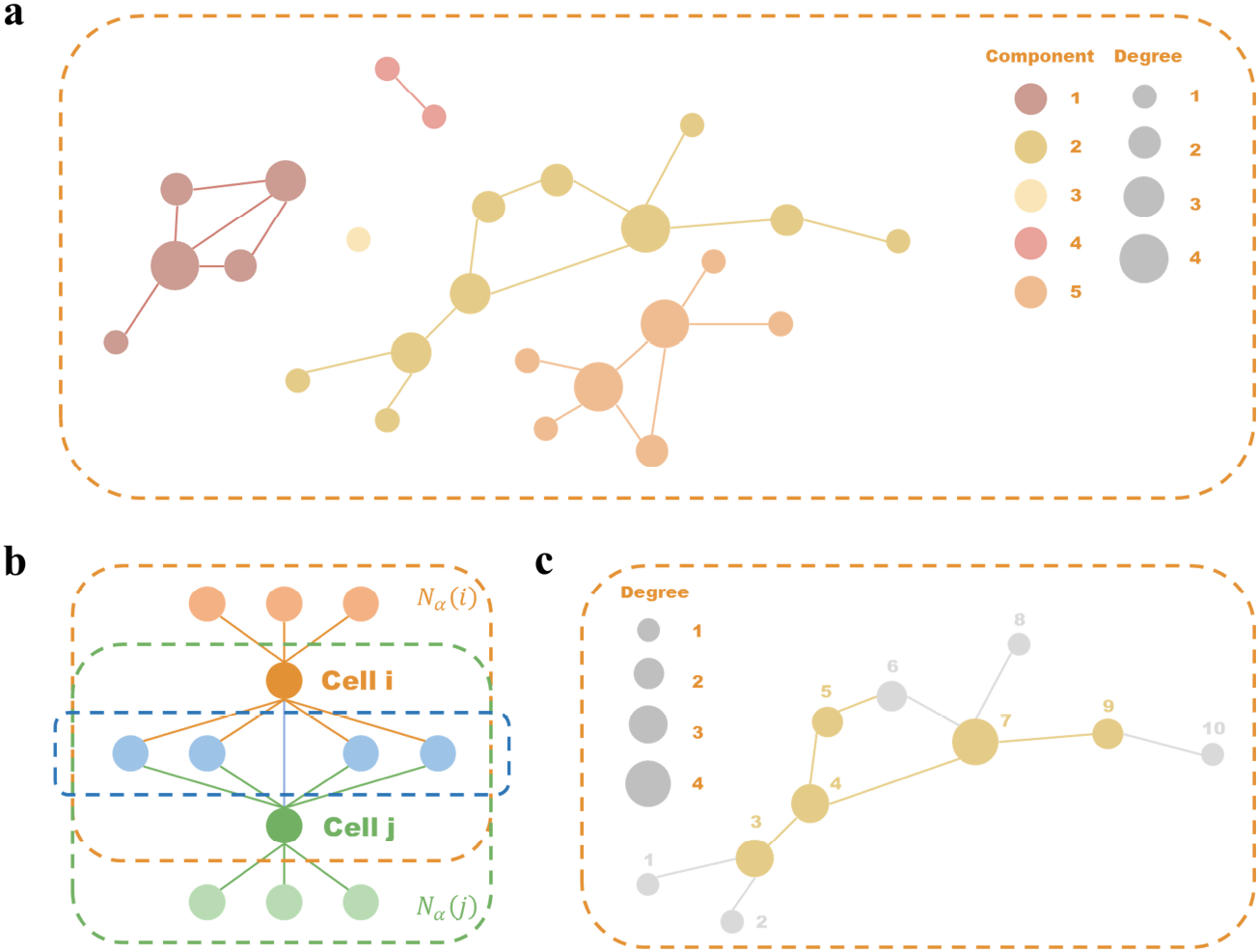
Construction of supercells. (a) An example of a graph 𝒢_*M*_ constructed with the matrix **M** as the adjacency matrix. (b) A diagram shows that cell *i* and cell *j* are second-order co-occurrence neighbors. The ones within the orange dashed box are the *α*-nearest neighbors of cell *i*, and those within the green dashed box are the *α*-nearest neighbors of cell *j*. If cell *j* is within *α*-nearest neighbors of cell *i*, and there are more than *β* common cells within their respective *α*-nearest neighbors as shown by the blue dashed box, then they are second-order co-occurrence neighbors. (c) An example of cells being eliminated with different probabilities in a certain connected component. Suppose we have 10 uniform distribution test results on [0, 1], which are 0.4677, 0.9383, 0.5542, 0.0975, 0.9667, 0.9137, 0.5838, 0.0260, 0.8815, and 0.8582. Then, from the calculation of Eq.(12) - Eq.(14), it can be known that cells 1, cell 2, cell 6, cell 8, and cell 10 are eliminated.

The cross-omics consistency matrix **M** = (*m*_*ij*_)_*n*×*n*_ ∈ {0, 1} ^*n*×*n*^ identifies persistent neighborhood relationships across modalities:

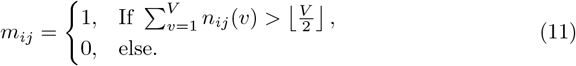

Using the consistency matrix **M** as an adjacency matrix, we construct the multi-omics similarity graph 𝒢_*M*_ in Fig. 18(a). Let {𝒢_*M*_ (1), 𝒢_*M*_ (2), …, 𝒢_*M*_ (*L*)} denote the *L* connected components of 𝒢_*M*_. While the connectedness principle suggests clustering co-component cells together, boundary cells with weak connectedness to other intra-component cells require special treatment. Therefore, we define cell’s higher-order neighborhood as the full set of cells that belong to the same connected components. We compute each cell’s degree centrality and then probabilistically discard cells whose profiles deviate from the group consensus.

For each component 𝒢_*M*_ (*l*) (*l* ∈ [1, *L*]), compute degree centrality of cell *i*:

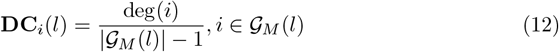

where |𝒢_*M*_ (*l*)| denotes component cardinality.

The high-order similarity metric of cell *i* in the *l*-th connected component is then:

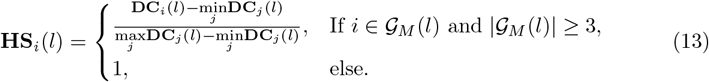

The cell elimination probability *P*_*i*_(*l*) follows:

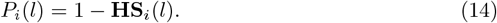

An iterative pruning process is implemented where cell *i* is removed if *U*_*i*_ ≤ *P*_*i*_(*l*) for *U*_*i*_ ∼ Uniform(0, 1). Fig. 18(c) demonstrates this workflow.

Each pruned component and eliminated cell is regarded as a “supercell”, which is a element in the set 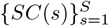

Then each cluster 𝒞_*p*_ becomes 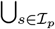 *SC*(*s*) where ℐ_*p*_ ⊆ {1, …, *S*}. The similarity between 𝒞_*p*_ and 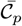 in the *v*-th omic is reformulated as:

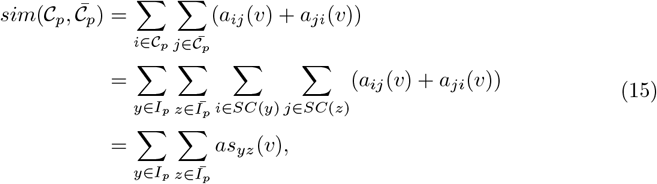

where the similarity between supercells is:

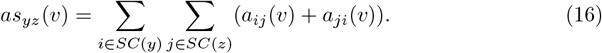

Then, the optimization framework transitions to:

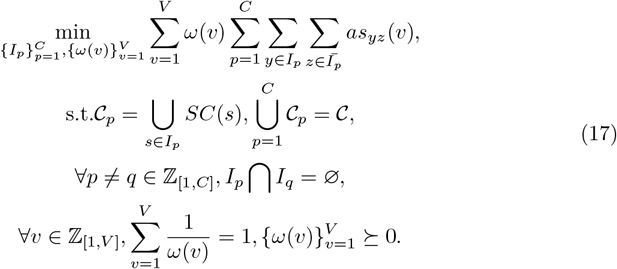

Further, the matrix formulation using cluster indicator matrix **E** ∈ {0, 1}^*S*×*C*^ is:

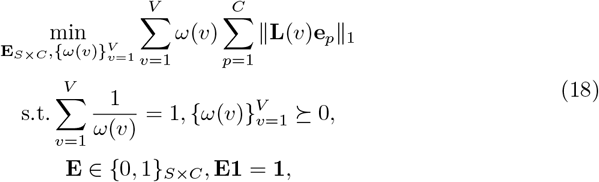

where **E**_*S*×*C*_ = (*e*_1_, *e*_2_, …, *e*_*C*_) represents the clustering result of supercells, **L**(*v*) is the Laplacian matrix of **AS**(*v*) = (*as*_*yz*_(*v*)) defined in Eq. (16).

We implement a block coordinate descent (BCD) optimization framework with alternating updates between cluster assignments **E**_*S*×*C*_ and modality weights 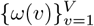. The iterative scheme proceeds as follows:

Firstly, given current cluster indicators **E**^(*t*)^, the analytical solution of 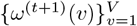 can be obtained as Eq. (19).

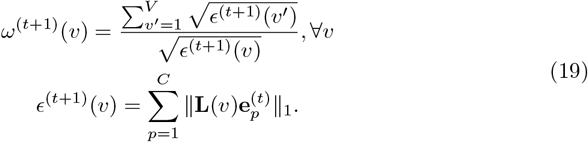

Secondly, fix 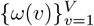 and update **E**_*S*×*C*_. We define the matrix 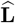 as Eq. (20).

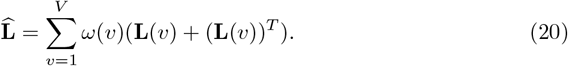

Meanwhile, define 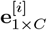 as the row vector where the *i*-th element is 1 and the remaining elements are 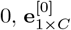 as a zero vector, and **E**^[**i**]^ as a matrix with **e**^[*i*]^ as the *d*-th row and the remaining rows being the same as **E**.

Next, we traverse the number of rows *d* and update the *d*-th row. Our objective function is:

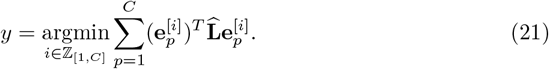

If we define:

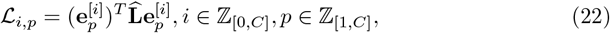

then the objective function is transformed into:

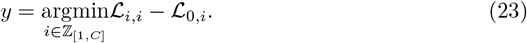

Find the index in the *d*-th row vector **e**^*d*^ where the element is 1 and denote it as *j*. Then, calculate ℒ _*i,i*_ − ℒ_0,*i*_ based on Eq. (24).

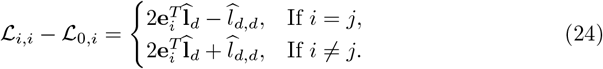

When traversing all ℒ_*i,i*_ − ℒ_0,*i*_ corresponding to *i*, the optimal solution *y* can be found, that is, the optimal solution in row *d*:

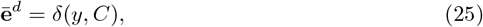

where δ(*y, C*) is a row vector where the *y*-th element is 1 and the remaining elements are 0.

Alternate update **E**_*S*×*C*_ and 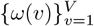 until convergence. The terminal cluster assignments are obtained by mapping **E** to supercell 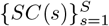.

#### Algorithm 1 The Algorithm of scHG

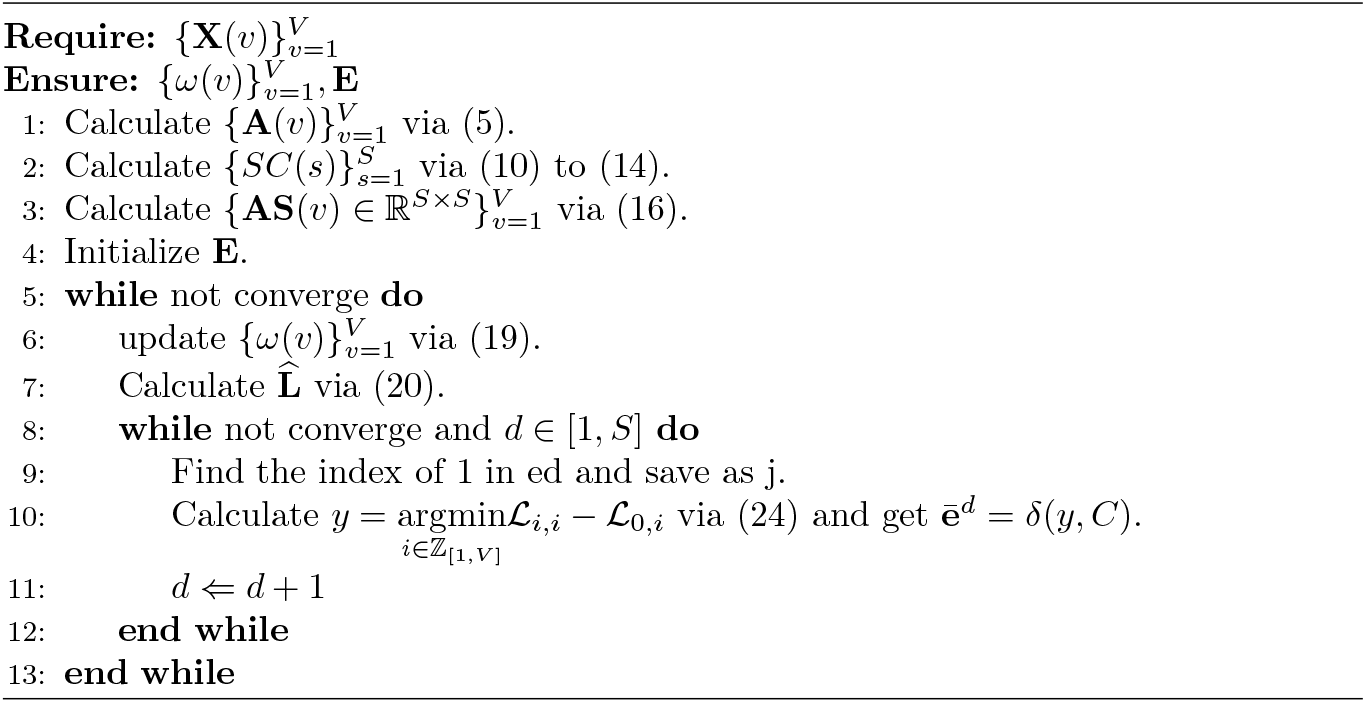

### Estimation of the cluster numbers

When the true cluster number is unknown, we propose a consensus estimation framework for multi-omics data 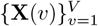. Each omic dataset is independently subjected to *v*-means clustering with elbow method optimization. For dataset **X**(*v*), the within-cluster sum of squared errors (SSE) for *C*(*v*) clusters is defined as:

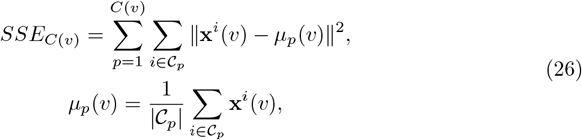

where 𝒞_*p*_ denotes the *p*-th cluster cell set.

The optimal cluster number *C*(*v*) per omic is determined by maximizing the elbow criterion:

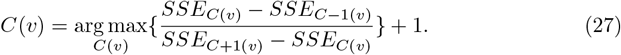

When omic-specific estimates 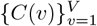 identical, the global cluster number *C* adopts their consensus value. For discrepancy cases, we compute the fused high-order multi-view similarity matrix **Ã** and its optimal *C*(**Ã**) using analogous criteria (Eq. (26)-(27)), then determine:

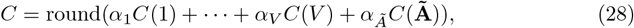

where *α*_*v*_ and *α*_**Ã**_ are weight coefficients.

### Time complexity analysis

Supercell constructing includes Eq. (10) to Eq. (14) executing in *O*(*n* + |*E*|) time, where *n* and |*E*| denote vertex and edge counts respectively. Supercell clustering optimization requires *O*(*S*^2^*V T*) operations, with *S* supercells, *V* omics, and *T* iterations. So the total complexity *O*(*n* + |*E*| + *S*^2^*V T*) simplifies to *O*(*n* + |*E*| + *S*^2^) given *V* ≪ *S*. Since |*E*| ≪ *n*^2^ and *S* ≪ *n* empirically, the effective complexity stays well below the quadratic upper bound *O*(*n*^2^).

### Datasets and preprocessing

We utilized five real-world datasets and one simulated dataset as shown in Table 7, which are elaborated as follows:

1. PBMC10× dataset: The 10× dataset contains 6661 cells with 7 cell types. The data were extracted from [54].
2. PBMC Inhouse dataset: The pbmc inhouse dataset contains 1182 cells with 6 cell types. The data were extracted from [15].
3. Bmcite dataset: The bmcite dataset contains 30672 cells with 27 cell types. The data were extracted from [55].
4. mESC dataset: The mESC dataset contains 77 cells with 2 cell types. The data were extracted from [56]. For each omics, we select the 125 columns of features with the largest variance.
5. PBMC Cao dataset: The 9631 dataset contains 8185 cells with 9 cell types. The data were extracted from [57].
6. Sim dataset: The Sim dataset contains 500 cells with 4 cell types. The data were extracted from [13].

**Table 7.**
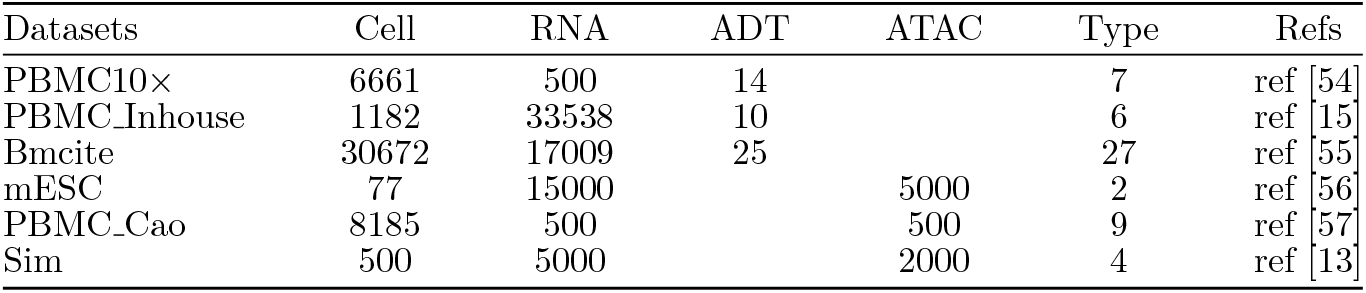
Detail information of single-cell multi-omics datasets.

| Datasets | Cell | RNA | ADT | ATAC | Type | Refs |
| --- | --- | --- | --- | --- | --- | --- |
| PBMC10× | 6661 | 500 | 14 | | 7 | ref [54] |
| PBMC_Inhouse | 1182 | 33538 | 10 |  | 6 | ref [15] |
| Bmcite | 30672 | 17009 | 25 |  | 27 | ref [55] |
| mESC | 77 | 15000 |  | 5000 | 2 | ref [56] |
| PBMC_Cao | 8185 | 500 |  | 500 | 9 | ref [57] |
| Sim | 500 | 5000 |  | 2000 | 4 | ref [13] |

For datasets where any omics modality contains ≥ 5000 features, we perform variance-based feature selection to retain the top 125 highest-variance features. This preprocessing ensures computational efficiency while preserving biologically informative signals.

### Compared methods

#### GBS [58]

GBS is a graph-based approach for Multi-View Clustering by proposing a unified framework studying generalization and graph metric impact. Its novel method effectively constructs adaptive graph matrices, automatically weights them, and directly produces final clusters.

#### scAI [13]

scAI is a single-cell aggregation and integration method designed to deconvolute cellular heterogeneity from parallel transcriptomic and epigenomic profiles by iteratively learning and aggregating sparse epigenomic signals across similar cells in an unsupervised manner, enabling coherent fusion with transcriptomics to dissect multi-omic heterogeneity and uncover regulatory mechanisms.

#### SMSC [59]

SMSC unifies joint nonnegative-spectral embedding with two distinctive features: (1) nonnegative embeddings directly yield cluster assignments, eliminating post-processing steps; (2) automatic parameter learning removes manual tuning requirements.

#### CGD [20]

CGD proposes a parameter-free, convergence-guaranteed approach for multi-view clustering by cross-view graph diffusion. It addresses key limitations of existing methods—model dependency, high computational complexity, and hyperparameter sensitivity—through an iterative diffusion process that 1) refines single-view graphs by preserving manifold structures and 2) leverages complementary information across views, yielding a unified graph that significantly outperforms benchmarks on seven evaluation metrics.

#### scMNMF [17]

scMNMF is an algorithm that jointly performs dimensionality reduction and cell clustering through non-negative matrix factorization (NMF) on single-cell multi-omics data.

#### GSTRPCA [60]

GSTRPCA is an adaptive tensor decomposition framework extending Tensor Robust Principal Component Analysis (TRPCA). It incorporates low-rank and sparse constraints via a weighted thresholding scheme, preserving data structure integrity while extracting latent cross-omic features.

### Hyperparameter configuration

The similarity matrix **A**(*v*) (Eq. (5)) employs adaptive regularization:

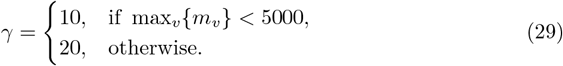

For second-order co-occurrence matrix **N**(*v*) (Eq. (10)), we fix *α* = 5 and *β* = 2. Cluster number estimation (Eq. (28)) uses modality-weighted fusion:

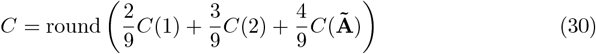

where weights 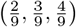 reflect the relative importance of RNA, ADT(or ATAC), and fused similarity features respectively.

### Performance metrics

Two established metrics are employed to assess clustering quality: Adjusted Rand Index (**ARI**) and Normalized Mutual Information (**NMI**). Consider two partitions: the ground truth partition 𝒯 = {𝒯_1_, 𝒯_2_, …, 𝒯_*t*_} and the clustered partition 𝒞= {𝒞_1_, 𝒞_2_, …, 𝒞_*p*_, where *t* and *p* denote the cardinality of true and clustered categories, respectively.

The **ARI** is formulated as:

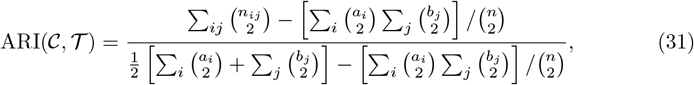

where *n*_*ij*_ denotes the overlap between 𝒯_*i*_ and 𝒞_*j*_, *a*_*i*_ = |𝒯_*i*_|, *b*_*j*_ = |𝒞_*j*_ |, and *n* is the total sample size. **ARI** ranges [-1,1], where 1 indicates a perfect match for the reference label.

The **NMI** is defined as:

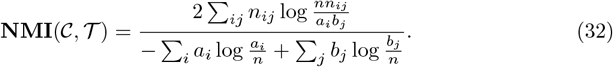

**NMI** ranges [0,1], where 1 indicates full information sharing.

Note that our ablation analysis further incorporated six validation metrics: Accuracy (**ACC**; [47]), **Purity** ([50]), **F1-score** ([51]), **Precision** ([51]), **Recall** ([51]), and Rand Index (**RI**; [52]).

## Data and code availability

All code and data for scHG are available at https://github.com/anchor-hue/scHG and http://mialab.ruc.edu.cn/scHG_code/zip.

## Acknowledgments

This work is supported in part by National Key Research and Development Program (2025YFC

2311702); National Natural Science Foundation of China (12426303 to X.Q.G.); Shenzhen Medical Research Fund (B2402038); Strategic Scientist Leadership Program (25XNKJ31 to X.Q.G.), Renmin University of China; Public Computing Cloud, Renmin University of China; Big Data and Responsible Artificial Intelligence for National Governance, Renmin University of China.

The analysis result of GO enrichment, GSEA analysis were generated using the R software packages “clusterProfiler” througn the CNSknowall (https://cnsknowall.com/index.html#/Home), a comprehensive web service for biomedical data analysis and visualization. We thank the CNSknowall platform (https://cnsknowall.com) for providing data analysis services.

## Author contributions

Y.X.H. and Y.G. conceived the study, designed the experiments, processed the data, analyzed the results, and wrote the manuscript. X.Q.G. supervised the study, provided conceptual guidance, and revised the manuscript.

## Footnotes

1 All experiments were executed on a Lenovo Legion R7000 2020 laptop equipped with an AMD Ryzen 7 4800H processor and 16 GB 3200 MT/s RAM, using MATLAB R2022a.

2 Null hypothesis H_0_: *μ*Target Cluster. ≤*μ*Others; alternative hypothesis *H*_1_: *μ*Target Cluster > *μ*Others.

3 GSTRPCA is inapplicable to datasets mESC and PBMC Cao, and SMSC yields a 1-dimensional latent space for this dataset; consequently, both methods lack t-SNE visualizations in the dataset mentioned above.

